# Mechanistic Analysis of the Broad Antiretroviral Resistance Conferred by HIV-1 Envelope Glycoprotein Mutations

**DOI:** 10.1101/2020.08.06.239350

**Authors:** Yuta Hikichi, Rachel Van Duyne, Phuong Pham, Jennifer L. Groebner, Ann Wiegand, John W. Mellors, Mary F. Kearney, Eric O. Freed

**Affiliations:** Virus-Cell Interaction Section, HIV Dynamics and Replication Program, NCI-Frederick, Frederick, Maryland, United States of America; Translational Research Section, HIV Dynamics and Replication Program, NCI-Frederick, Frederick, Maryland, United States of America; Department of Immunology, University of Pittsburgh, Pittsburgh, Pennsylvania, United States of America

**Author notes:** Corresponding author (EOF). Department of Pharmacology and Physiology, College of Medicine, Drexel University, Philadelphia, Pennsylvania, United States of America (Rachel Van Duyne).

## Abstract

Despite the effectiveness of antiretroviral (ARV) therapy, virological failure can occur in some HIV-1 infected patients in the absence of mutations in the proteins targeted by these drugs. We previously reported that, *in vitro*, the lab-adapted NL4-3 strain of HIV-1 can acquire resistance to the integrase inhibitor dolutegravir (DTG) by acquiring mutations in the envelope glycoprotein (Env) that enhance the ability of HIV-1 to spread via cell-cell transmission. In this study, we investigated whether Env-mediated drug resistance extends to ARVs other than DTG and whether it occurs in other HIV-1 isolates. We demonstrate that Env mutations can broadly confer resistance to multiple classes of ARVs in the context of cell-cell but not cell-free infection and also increase resistance to ARVs when coupled with target-gene drug resistance mutations. To investigate the mechanism of Env-mediated drug resistance, we evaluated the impact of the Env mutations on Env stability and conformational dynamics. We observe that the NL4-3 Env mutants display a more stable and closed Env conformation compared to WT virus and reduced rates of gp120 shedding. We also selected for mutations in the gp41 ectodomain of clinically relevant, CCR5-tropic isolates in the presence of DTG. These Env mutants exhibit reduced susceptibility to DTG, with effects on replication kinetics and Env structure that are HIV-1 strain-dependent. Finally, to examine a possible *in vivo* relevance of Env-mediated drug resistance, we performed single-genome sequencing of plasma-derived virus from five patients failing an integrase inhibitor-containing regimen. This analysis revealed the presence of several mutations in the highly conserved gp120-gp41 interface despite low frequency of resistance mutations in integrase. These results suggest a “stepping-stone” model whereby mutations in Env that enhance the ability of HIV-1 to spread via a cell-cell route increase the opportunity for the virus to acquire high-level drug resistance mutations in ARV-target genes.

**Author summary:** Although combination antiretroviral (ARV) therapy has proven highly effective in controlling the progression of HIV disease, drug resistance can be a major obstacle to long-term treatment, particularly in resource-limited settings. In most cases, resistance arises from the accumulation of mutations in the ARV-target genes; however, in some cases, resistance develops without ARV target-gene mutations. We previously reported that mutations in the HIV-1 envelope glycoprotein (Env) confer resistance to an integrase inhibitor. Here we investigated the mechanism of Env-mediated drug resistance and the possible contribution of Env to virological failure *in vivo*. We demonstrate that Env mutations can confer broad resistance to multiple classes of ARVs and define the effect of the Env mutations on Env subunit interactions and sensitivity to neutralizing antibodies. We also selected for drug resistance mutations in Env in clinically relevant HIV-1 isolates. We observed that many Env mutations accumulated in individuals failing integrase inhibitor therapy despite a low frequency of resistance mutations in integrase. Our findings suggest that broad-based, Env-mediated drug resistance may impact current and possibly future therapeutic strategies. Our findings also provide clues towards understanding how ARV-treated patients can experience virological failure without acquiring drug resistance mutations in ARV-target genes.

## Introduction

More than thirty antiretrovirals (ARVs) are currently approved to treat HIV-1 infection [1]. These ARVs are categorized into five different classes based on their targets: (1) nucleoside analog reverse transcriptase inhibitors (NRTIs), (2) non-nucleoside analog reverse transcriptase inhibitors (NNRTIs), (3) integrase (IN) strand transfer inhibitors (INSTIs), (4) protease (PR) inhibitors (PIs) and (5) entry inhibitors (Ent-Is). The use of combinations of these ARVs (combination antiretroviral therapy; cART) has proven remarkably effective in controlling HIV disease progression and prolonging survival. However, resistance to ARVs emerges in some patients because of poor adherence, use of a suboptimal drug regimen, and/or lack of viral load monitoring, particularly in resource-limited settings. Transmitted drug resistance is therefore becoming an increasingly serious problem in many parts of the world [2]. In most cases, HIV drug resistance is the consequence of mutations that emerge in the viral genes targeted by the drugs, and the molecular and structural basis for such resistance has been extensively studied [2]. However, particularly in the case of PIs and INSTIs, virological failure can occur in the absence of target-gene mutations [3–10], indicating that mutations outside the target gene can contribute to drug resistance. For example, *in vitro* selection studies have demonstrated that mutations in the 3’ polypurine tract (3’ PPT) or in the viral long terminal repeat (LTR) can confer resistance to INSTIs [11, 12], and mutations in Gag and the envelope glycoprotein (Env) have been implicated in PI resistance [13, 14].

HIV-1 Env is translated as a gp160 precursor, which is cleaved by a cellular furin-like protease to generate noncovalently linked gp120 and gp41 subunits that are incorporated into virus particles as a heterotrimeric spike [15]. The binding of gp120 to CD4 on the target cell surface triggers a conformational rearrangement in Env that exposes the coreceptor (CCR5 or CXCR4)-binding site. Interaction of gp120 with coreceptor promotes the refolding of gp41 heptad repeat 1 and 2 (HR1 and 2) to form an antiparallel six-helix bundle that mediates the fusion of viral and cellular membranes, allowing viral entry into the cytosol of the target cell [16].

HIV-1 Env is the only viral protein exposed on the surface of the infected cell or viral particle; it is therefore the target of neutralizing antibodies (NAbs) that can block viral entry and induce antibody-mediated effector functions, such as antibody-dependent cellular cytotoxicity (ADCC). NAbs recognize conserved regions of the Env trimer, including (1) the CD4-binding site (CD4bs), (2) V2 apex, (3) V3 glycan, (4) gp120-gp41 interface, and (5) gp41 membrane-proximal external region (MPER) [17]. Single-molecule fluorescence resonance energy transfer (smFRET) analysis has revealed that Env trimers fluctuate between closed (state 1), intermediate (state 2) and open (state 3) conformations [18, 19]. The binding properties of NAbs are influenced by the conformational state of Env. For example, while many anti-V2 apex and anti-CD4bs Abs preferentially recognize the closed Env conformation, antibodies that bind the gp41 MPER and a CD4-induced epitope (CD4i) preferentially recognize Env conformations that arise after the engagement of gp120 with CD4. Therefore, NAbs are useful as molecular probes to investigate the structure and conformational dynamics of Env [18, 19].

HIV-1 can spread either via a cell-free route or by cell-cell transmission at points of cell-cell contact known as virological synapses (VSs). The formation of a VS is initiated by the interaction of Env on the infected cell and CD4 on the target cell [20–22], although Gag can accumulate at the VS even in the absence of Env [23]. The VS is stabilized by cellular adhesion proteins, such as LFA-1 and ICAM-1, and lipid raft microdomains and the actin cytoskeleton are also implicated in VS formation [24–26]. At least *in vitro*, cell-cell transfer of HIV-1 is markedly more efficient than cell-free infection [27–29]. HIV-1 transmission at a VS provides a higher multiplicity of infection (MOI) than cell-free infection, resulting in multiple copies of proviral DNA in the target cells [30–33]. This high MOI allows HIV-1 to overcome multiple barriers to infection, such as those imposed by ARVs, NAbs, and inhibitory host factors [32–38]. Although it is challenging to directly compare the relative contribution of cell-free infection and cell-cell transfer to HIV-1 dissemination and pathogenesis *in vivo*, several studies have suggested the importance of cell-associated virus in HIV-1 propagation in animal models of HIV-1 infection. Intravital imaging using humanized mice has demonstrated that HIV-1-infected T-cells are motile in lymph nodes and spleen, and form transient, Env-dependent adhesive contacts that may facilitate cell-cell viral transfer [39, 40]. In addition, multicopy transmission of HIV-1 as foci within lymphoid tissue has been reported in humanized mice [30, 40]. Inhibition of egress of lymphocytes reduced systemic dissemination of HIV-1, suggesting that migration of infected cells plays an important role in virus spread to other organs [39]. Electron tomography and immunofluorescence analyses in gut lymphoid tissues and bone marrow of humanized mice have provided support for both cell-free and cell-cell modes of HIV-1 transmission [41–43]. If cell-cell transfer, and the resulting higher MOI, is a common occurrence *in vivo* one would expect to find large numbers of multiply infected cells in HIV-1-infected individuals. This has been observed in some [44–46] but not other [47] studies. Further analysis will be needed to understand the importance of cell-cell transfer in viral dissemination *in vivo*.

We previously reported that HIV-1 evades blocks to virus replication by acquiring Env mutations that enhance the capacity of the virus to spread via cell-cell transmission [48]. By propagating HIV-1 mutants containing substitutions in the p6 domain of Gag that severely delay virus replication [49], we selected for second-site compensatory mutations Env-Y61H and P81S in gp120 and the Env-A556T mutation in gp41. These Env mutations decrease the sensitivity of the virus to the highly potent INSTI dolutegravir (DTG). Moreover, propagation of HIV-1 in the presence of DTG led to the selection of the DTG-escape mutant Env-A539V in the absence of any mutations in IN. These four Env mutations cluster within the C1 domain of gp120 and the HR1 domain of gp41 (Fig 1), which have been shown through mutagenesis and structural studies to be critical for the stability of gp120-gp41 association in the unliganded (non-CD4-bound) state [50–52]. These results suggest that Env mutations that increase cell-cell spread may alter the stability of the gp120-gp41 interaction and demonstrate that mutations in Env can confer resistance to an ARV, at least in a subtype B, CXCR4-tropic, laboratory-adapted HIV-1 strain. Whether Env-mediated drug resistance occurs in clinical HIV-1 isolates is unclear because Env from laboratory-adapted and primary HIV-1 strains differs in significant functional and structural aspects. Laboratory-adapted isolates often sample the open Env conformation and are more sensitive to NAbs compared to primary isolates, as *in vitro* adaptation of HIV-1 in the absence of immune selective pressure often leads to the emergence of neutralization-sensitive variants [18, 53, 54]. In addition, laboratory-adapted isolates are more sensitive to soluble CD4 (sCD4) than primary isolates. sCD4 inhibits HIV-1 entry not only by competing with CD4 on the target cell, but also by inducing the shedding of gp120 from the surface of viral particles and infected cells. Env complexes from laboratory-adapted isolates are more prone to sCD4-induced gp120 shedding compared to primary isolates [55–57], indicating that Env stability differs between laboratory-adapted and primary isolates.

**Fig 1.**
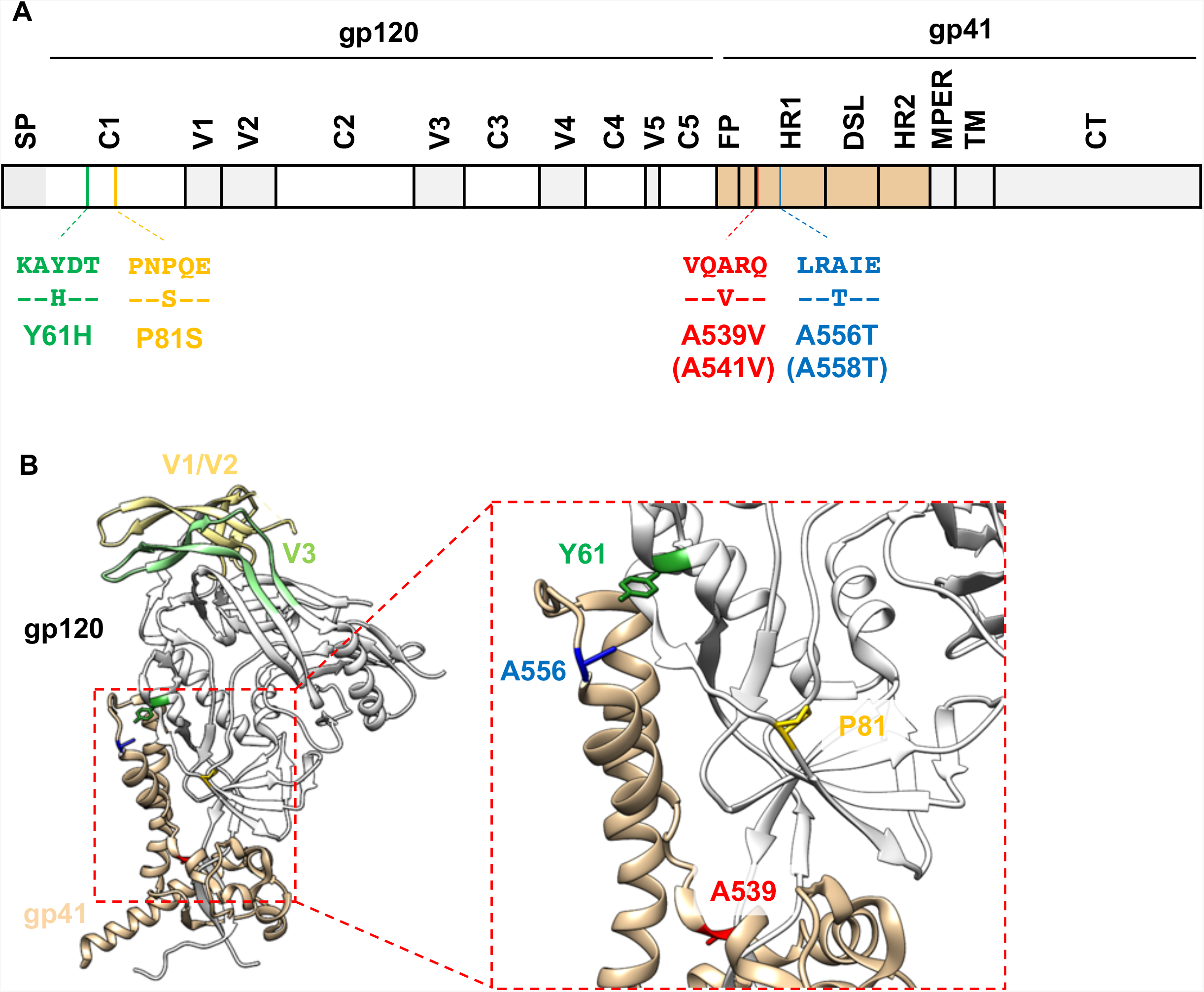
The location of previously reported Env mutations that overcome blocks to virus replication (Van Duyne et al., PNAS, 2019 [48]) (A) Schematic of the HIV-1 Env-coding region with the position of mutations indicated using NL4-3 (and HXB2, in parentheses) numbering. Labeled domains are defined as follows: C1–C5, constant region 1–5; V1–V5, variable region 1–5; FP, fusion peptide; HR1/HR2, heptad repeat 1/2; DSL, disulfide loop; MPER, membrane proximal external region; TM, transmembrane; CT, cytoplasmic tail. (B) A prefusion Env structure of subtype B JR-FL SOSIP.664 (PDB accession number 5FYK [96]) with the position of the Env mutations highlighted. Most of gp120 is shown in white, with gp120 V1/V2 and V3 loops colored in light yellow and light green, respectively. gp41 is colored in tan. Y61, P81, A539 and A556 are highlighted in green, yellow, red and blue, respectively. Structural model was generated using UCSF Chimera software [129].

In the present study, we examined whether our previously described Env mutations confer resistance not only to DTG but also to other classes of ARVs. To evaluate the impact of the Env mutations on the conformation and stability of the Env complex, we examined their effects on sensitivity to NAbs and on gp120 shedding. To further characterize the mechanism of Env-mediated drug resistance, we obtained Env mutants through *in vitro* selection experiments by propagating clinically relevant HIV-1 clones in the presence of DTG and analyzed the phenotypes of the selected Env mutants. Finally, to address the possible *in vivo* relevance of Env-mediated drug resistance, we used single-genome sequencing to compare viral sequences obtained from the plasma of individuals failing a raltegravir (RAL)-containing regimen. The results indicate that Env mutations confer resistance across a broad range of ARVs *in vitro*, and that the NL4-3 Env mutations decrease gp120 shedding and alter the sensitivity of Env to NAbs. Finally, we show that in some individuals failing a RAL-containing regimen in the absence of resistance mutations in IN, changes arise at the highly conserved gp120-gp41 interface, as observed during INSTI escape *in vitro*.

## Results

### The A539V mutation in gp41 provides a replication advantage over WT NL4-3 in the presence of multiple classes of ARVs

We previously reported that the Env-A539V mutation provides resistance to the INSTI DTG [48]. To examine whether Env-A539V also provides a selective replication advantage in the presence of other ARVs, the SupT1 T-cell line was transfected with the WT NL4-3 molecular clone or the Env-A539V derivative in the absence or presence of various concentrations (0.1 – 3,000 nM) of a panel of ARVs. Replication kinetics were monitored by measuring the RT activity in the cell culture media. Consistent with our previous report [48], Env-A539V exhibited faster replication kinetics compared to WT in the absence of drugs (Fig 2). Whereas replication of WT NL4-3 was largely inhibited in the presence of 3.0 nM DTG, Env-A539V could replicate, albeit with delayed kinetics (Fig 2A). By calculating the DTG IC_50_ based on the RT activity at the peak of replication in the absence of drugs, we found that Env-A539V showed 8.1-fold resistance to DTG relative to WT (Fig 2I). We also examined the sensitivity of Env-A539V to another INSTI, raltegravir (RAL). Env-A539V could replicate in the presence of 10 nM RAL and showed 5.4-fold resistance relative to WT NL4-3 (Figs 2B and I). We also measured replication kinetics of Env-A539V in the presence of the NNRTI efavirenz (EFV) and the NRTI emtricitabine (FTC). Whereas replication of WT NL4-3 was strongly inhibited in the presence of 3.0 nM EFV or 30 nM FTC, Env-A539V could still replicate at these concentrations (Figs 2C and D). IC_50_ calculations showed that Env-A539V exhibited 28- and 5.4-fold resistance against EFV and FTC, respectively (Fig 2I). Next, we examined the impact of PIs on the replication kinetics of Env-A539V. While replication of WT NL4-3 was strongly inhibited in the presence of 3.0 nM nelfinavir (NFV), the Env-A539V mutant could replicate and exhibited 24-fold resistance (Figs 2E and I). We also examined the sensitivity of Env-A539V to the allosteric integrase inhibitor (ALLINI), BI-224436, which induces aberrant IN multimerization and impairs the interaction between IN and the cellular cofactor LEDGF/p75 [58]. As shown in Fig 2F, Env-A539V caused a small but statistically significant reduction in the sensitivity to BI-224436 relative to WT NL4-3 (2.3-fold resistance, Fig 2I). These data demonstrate that Env mutations can overcome inhibition imposed by multiple classes of ARVs targeting not only post-entry steps, but also the maturation step of the HIV-1 replication cycle.

**Fig 2.**
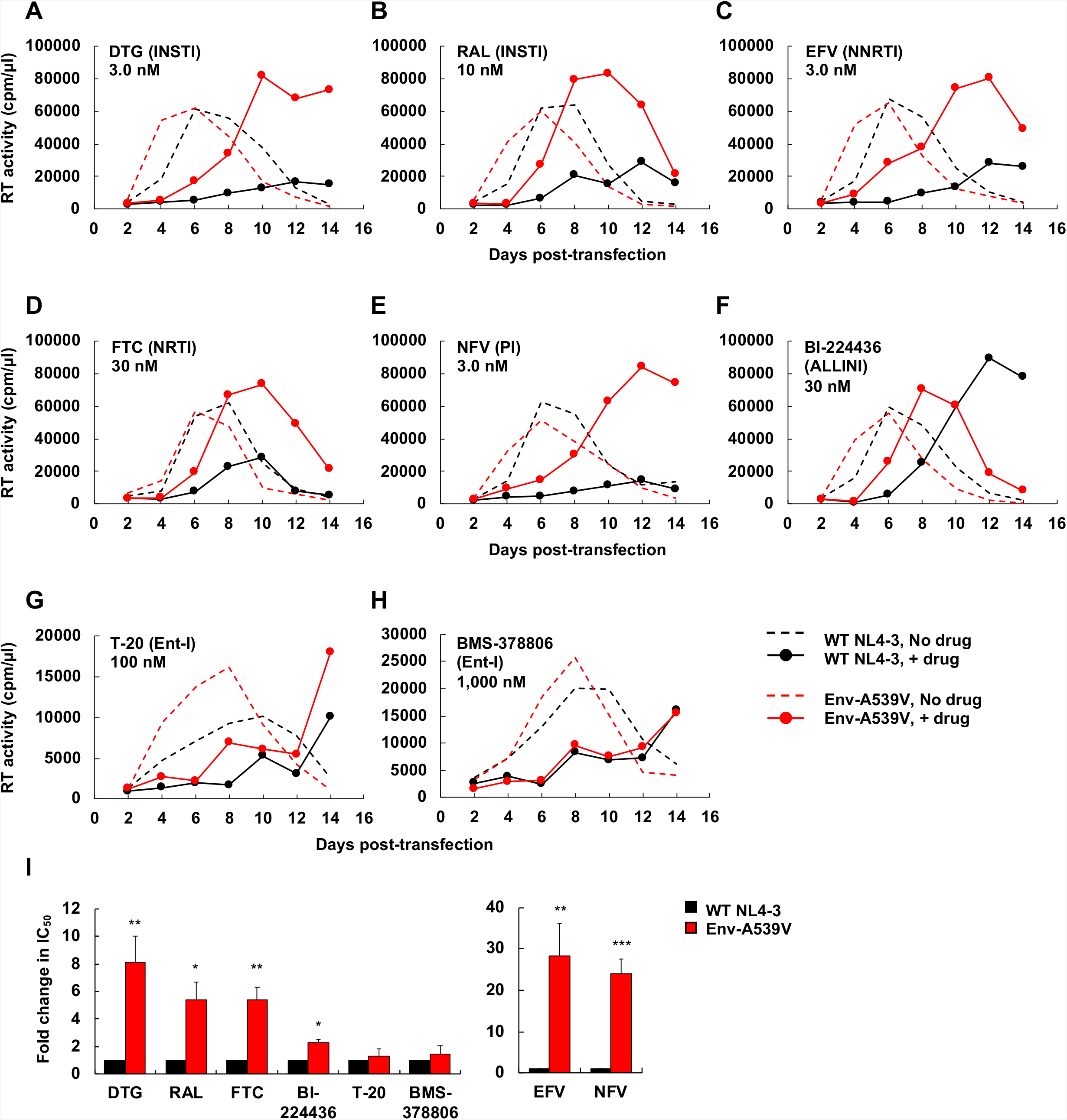
Replication kinetics of Env-A539V in the presence of INSTIs, NRTI, NNRTI, PI, ALLINI and Ent-Is. The SupT1 T-cell line was transfected with WT or Env-A539V proviral clones in the absence or in the presence of indicated concentrations of ARVs. (A and B) INSTIs, (C and D) NNRTI and NRTI, (E) PI, (F) ALLINI, and (G and H) Ent-Is. Virus replication kinetics were monitored by measuring RT activity at the indicated time points. Data are representative of at least two independent experiments. (I) The SupT1 T-cell line was transfected with WT or Env-A539V proviral clones in the absence or in the presence of serial dilutions (0.01 – 3,000 nM) of ARVs. IC_50_ values were calculated based on RT levels at the peak of virus replication. Fold changes in IC_50_ were calculated compared to WT. Data from at least two independent experiments are shown as means ± SE. *p*-values < 0.001 (***), < 0.01 (**), < 0.05 (*) by unpaired *t*-test. The EFV and NFV data are shown on a separate bar graph to avoid compression of the y-axis.

It has been suggested that entry inhibitors (Ent-Is) are effective in the context of cell-cell transmission [59, 60]. In addition, several studies have shown that Env mutations selected in the presence of Ent-Is altered viral sensitivity to other anti-Env agents [61–64]. These findings raise the possibility that Env mutations selected in the presence of ARVs might alter the viral sensitivity to Env-targeted inhibitors. To examine this question, we measured replication kinetics in the presence of the fusion inhibitor T-20 and the attachment inhibitor BMS-378806. Interestingly, these Ent-Is could efficiently suppress replication of both WT NL4-3 and Env-A539V with no statistically significant differences in antiviral IC_50_ in a multi-cycle spreading infection (Figs 2G – I). These observations suggest that Ent-Is could suppress the replication of Env mutants selected under the pressure of ARVs targeting the viral enzymes.

### The Env-A539V mutation does not alter drug sensitivity in cell-free infection

We previously proposed that Env mutations that confer resistance to ARVs do so by enhancing the efficiency of cell-cell transfer [48]. Based on this hypothesis, the Env mutations would not be predicted to confer resistance in the context of cell-free infectivity. To test this hypothesis, we measured the infectivity of WT NL4-3 and the Env-A539V mutant in the TZM-bl indicator cell line. For early-acting inhibitors (INSTI, NNRTI, NRTI and Ent-Is), infection of TZM-bl cells was performed in the presence of the drugs. For late-acting inhibitors (PI and ALLINIs), virus stocks were produced in the presence of the inhibitors, and then used to infect the TZM-bl cells. As we reported previously [48], Env-A539V showed comparable cell-free infectivity relative to WT NL4-3 in the absence of inhibitor (Fig 3A and Table 1). Consistent with the hypothesis that the reduced ARV sensitivity of the Env-A539V mutant is conferred at the level of cell-cell transmission, WT and Env-A539V infectivity was reduced to the same extent by the inhibitors (Figs 3B – I, and Table 1). Interestingly, Env-A539V did not alter susceptibility to either T-20 or BMS-378806, suggesting that Ent-Is efficiently suppress both cell-free infection and cell-cell transmission of HIV-1.

**Fig 3.**
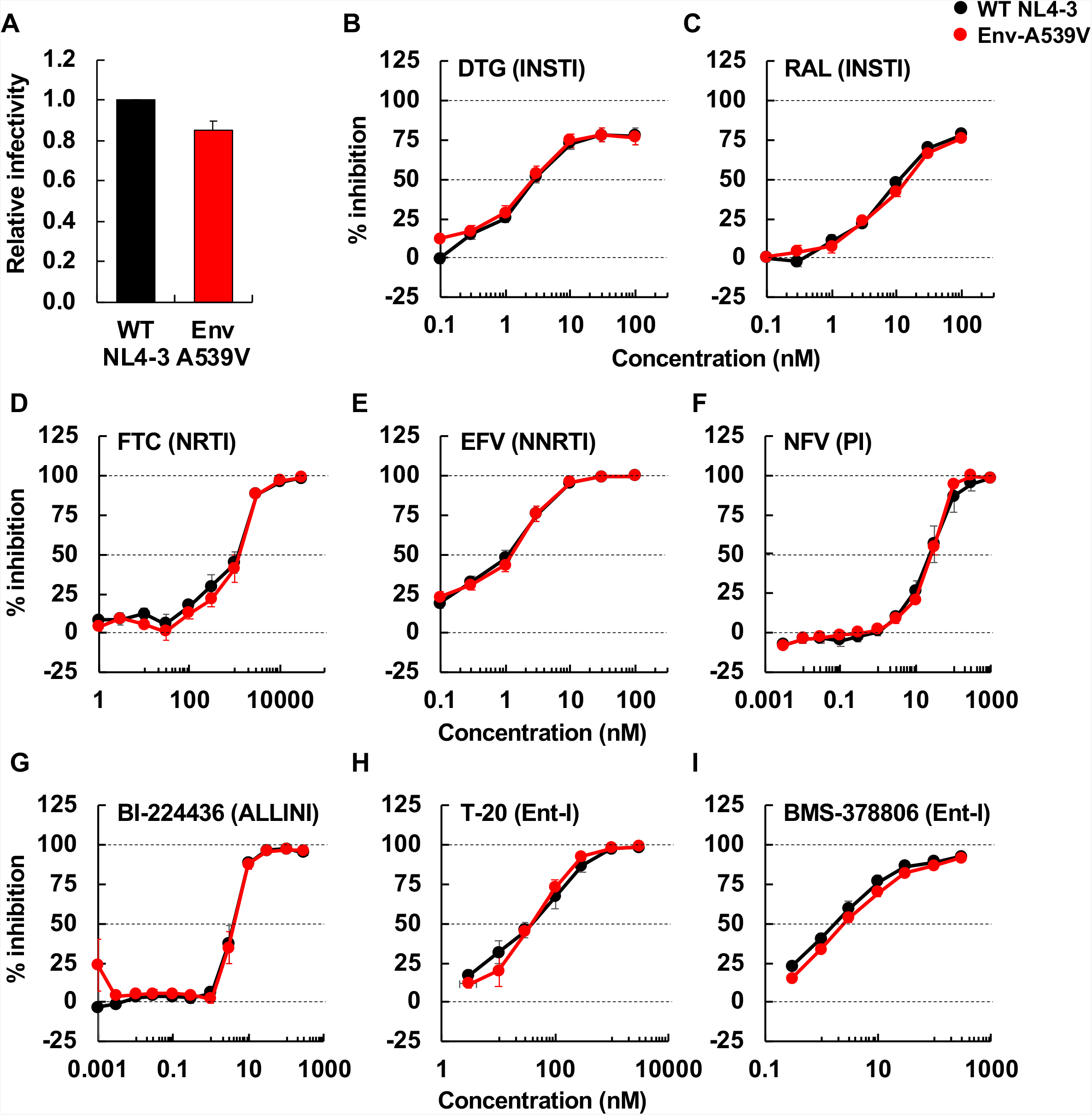
Cell-free infectivity of Env-A539V in the presence or absence of ARVs. (A) RT-normalized virus stocks produced from HeLa cells were used to infect TZM-bl cells. Luciferase activity was measured at 48 hrs post-infection. Infectivity of WT NL4-3 is normalized to 1.0. (B – I) TZM-bl cells were exposed to 100 TCID_50_ of WT or Env-A539V virus in the presence of varying concentrations (from 0.003 – 3,000 nM) of ARVs and incubated for 48 hrs. For NFV and BI-224436, 293T cells were transfected with the indicated proviral clones in the presence of the inhibitors. At 48 hrs post-transfection, the supernatants were used to infect TZM-bl cells. Data from at least three independent experiments are shown as mean ± SE.

**Table 1.**
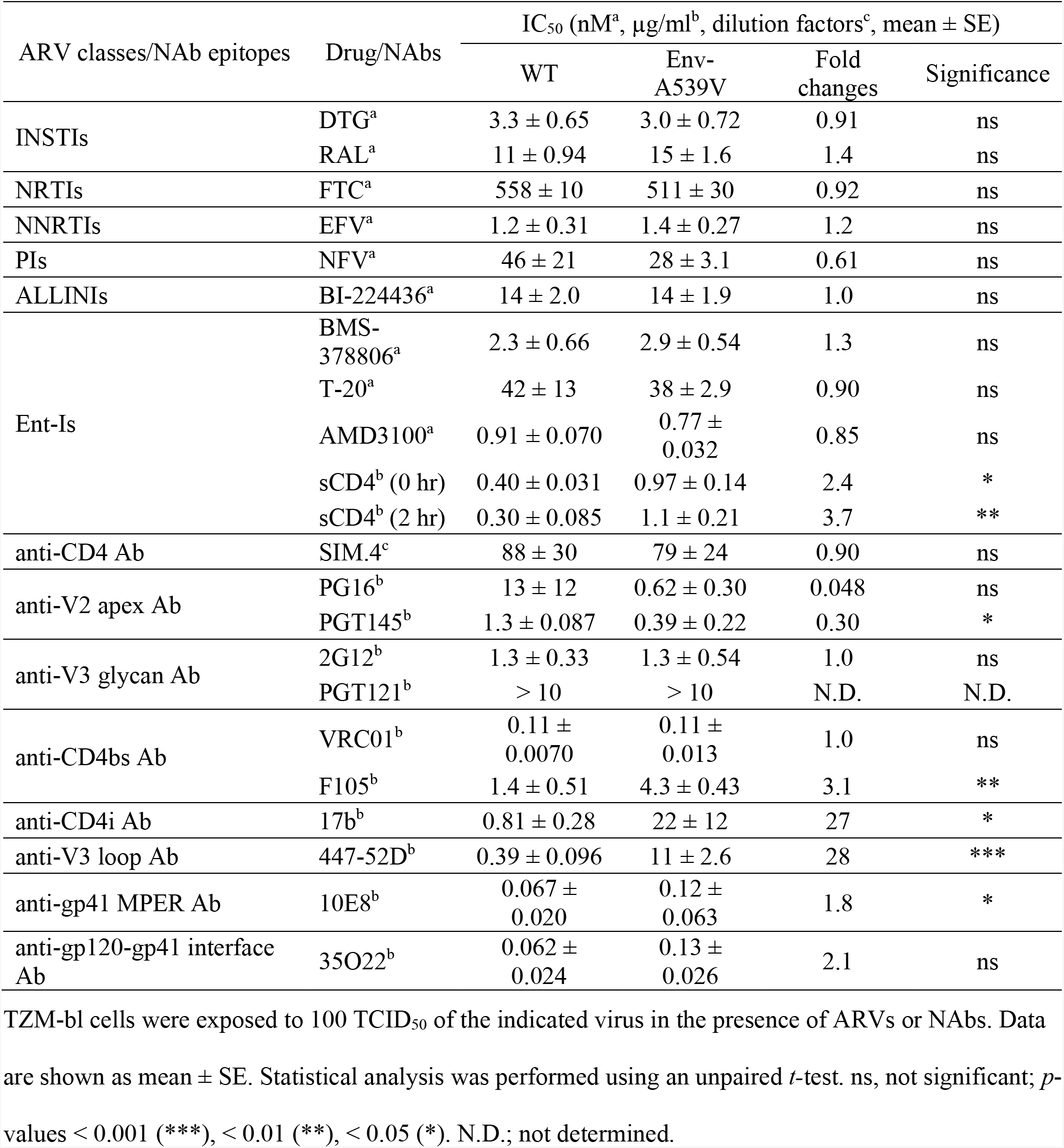
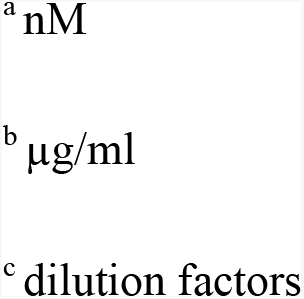
IC_50_ values in cell-free infection for WT NL4-3 and the A539V variant.

| ARV classes/NAb epitopes | Drug/NAbs | IC<sub>50</sub> (nM<sup>a</sup>, µg/ml<sup>b</sup>, dilution factors<sup>c</sup>, mean ± SE) |  |  |  |
| --- | --- | --- | --- | --- | --- |
|  |  | WT | Env-A539V | Fold changes | Significance |
| INSTIs | DTG<sup>a</sup> | 3.3 ± 0.65 | 3.0 ± 0.72 | 0.91 | ns |
|  | RAL<sup>a</sup> | 11 ± 0.94 | 15 ± 1.6 | 1.4 | ns |
| NRTIs | FTC<sup>a</sup> | 558 ± 10 | 511 ± 30 | 0.92 | ns |
| NNRTIs | EFV<sup>a</sup> | 1.2 ± 0.31 | 1.4 ± 0.27 | 1.2 | ns |
| PIs | NFV<sup>a</sup> | 46 ± 21 | 28 ± 3.1 | 0.61 | ns |
| ALLINIs | BI-224436<sup>a</sup> | 14 ± 2.0 | 14 ± 1.9 | 1.0 | ns |
| Ent-Is | BMS-378806<sup>a</sup> | 2.3 ± 0.66 | 2.9 ± 0.54 | 1.3 | ns |
|  | T-20<sup>a</sup> | 42 ± 13 | 38 ± 2.9 | 0.90 | ns |
|  | AMD3100<sup>a</sup> | 0.91 ± 0.070 | 0.77 ± 0.032 | 0.85 | ns |
|  | sCD4<sup>b</sup> (0 hr) | 0.40 ± 0.031 | 0.97 ± 0.14 | 2.4 | * |
|  | sCD4<sup>b</sup> (2 hr) | 0.30 ± 0.085 | 1.1 ± 0.21 | 3.7 | ** |
| anti-CD4 Ab | SIM.4<sup>c</sup> | 88 ± 30 | 79 ± 24 | 0.90 | ns |
| anti-V2 apex Ab | PG16<sup>b</sup> | 13 ± 12 | 0.62 ± 0.30 | 0.048 | ns |
|  | PGT145<sup>b</sup> | 1.3 ± 0.087 | 0.39 ± 0.22 | 0.30 | * |
| anti-V3 glycan Ab | 2G12<sup>b</sup> | 1.3 ± 0.33 | 1.3 ± 0.54 | 1.0 | ns |
|  | PGT121<sup>b</sup> | > 10 | > 10 | N.D. | N.D. |
| anti-CD4bs Ab | VRC01<sup>b</sup> | 0.11 ± 0.0070 | 0.11 ± 0.013 | 1.0 | ns |
|  | F105<sup>b</sup> | 1.4 ± 0.51 | 4.3 ± 0.43 | 3.1 | ** |
| anti-CD4i Ab | 17b<sup>b</sup> | 0.81 ± 0.28 | 22 ± 12 | 27 | * |
| anti-V3 loop Ab | 447-52D<sup>b</sup> | 0.39 ± 0.096 | 11 ± 2.6 | 28 | *** |
| anti-gp41 MPER Ab | 10E8<sup>b</sup> | 0.067 ± 0.020 | 0.12 ± 0.063 | 1.8 | * |
| anti-gp120-gp41 interface Ab | 35O22<sup>b</sup> | 0.062 ± 0.024 | 0.13 ± 0.026 | 2.1 | ns |

TZM-bl cells were exposed to 100 TCID<sub>50</sub> of the indicated virus in the presence of ARVs or NAbs. Data are shown as mean ± SE. Statistical analysis was performed using an unpaired *t*-test. ns, not significant; *p*-values < 0.001 (\*\*\*), < 0.01 (\*\*), < 0.05 (\*). N.D.; not determined.
<sup>a</sup> nM
<sup>b</sup> µg/ml
<sup>c</sup> dilution factors

### The Env-A539V mutation increases viral resistance to EFV when coupled with the RT-Y188L mutation

To examine whether mutations in Env can increase the level of resistance conferred by drug target gene mutations, we tested EFV resistance of the RT-Y188L mutant [65] in the context of either WT or A539V Env. As shown in Fig 4A (left panel), RT-Y188L showed a modest delay in replication compared to WT in the absence of EFV, indicating that this RT mutation confers a small fitness defect. This replication defect was rescued by Env-A539V, consistent with our earlier finding that Env mutations can rescue the replication defect conferred by a mutation in IN [48]. In the presence of 1.0 mM EFV, the Env-A539V mutation was able to markedly enhance the replication kinetics of the RT-Y188L mutant (Fig 4A, right panel), a result that was confirmed over a broad range of EFV concentrations (Figs 4B and D). In contrast to the results in the multi-cycle spreading infection (Fig 4B), the Env-A539V mutation did not increase EFV resistance of RT-Y188L in a single-cycle infectivity assay (Figs 4C and D). Taken together, these results demonstrate that, in the context of a spreading infection in which virus transmission can take place via cell-cell transfer, an Env mutation can significantly increase the resistance conferred by a mutation in an ARV-target gene.

**Fig 4.**
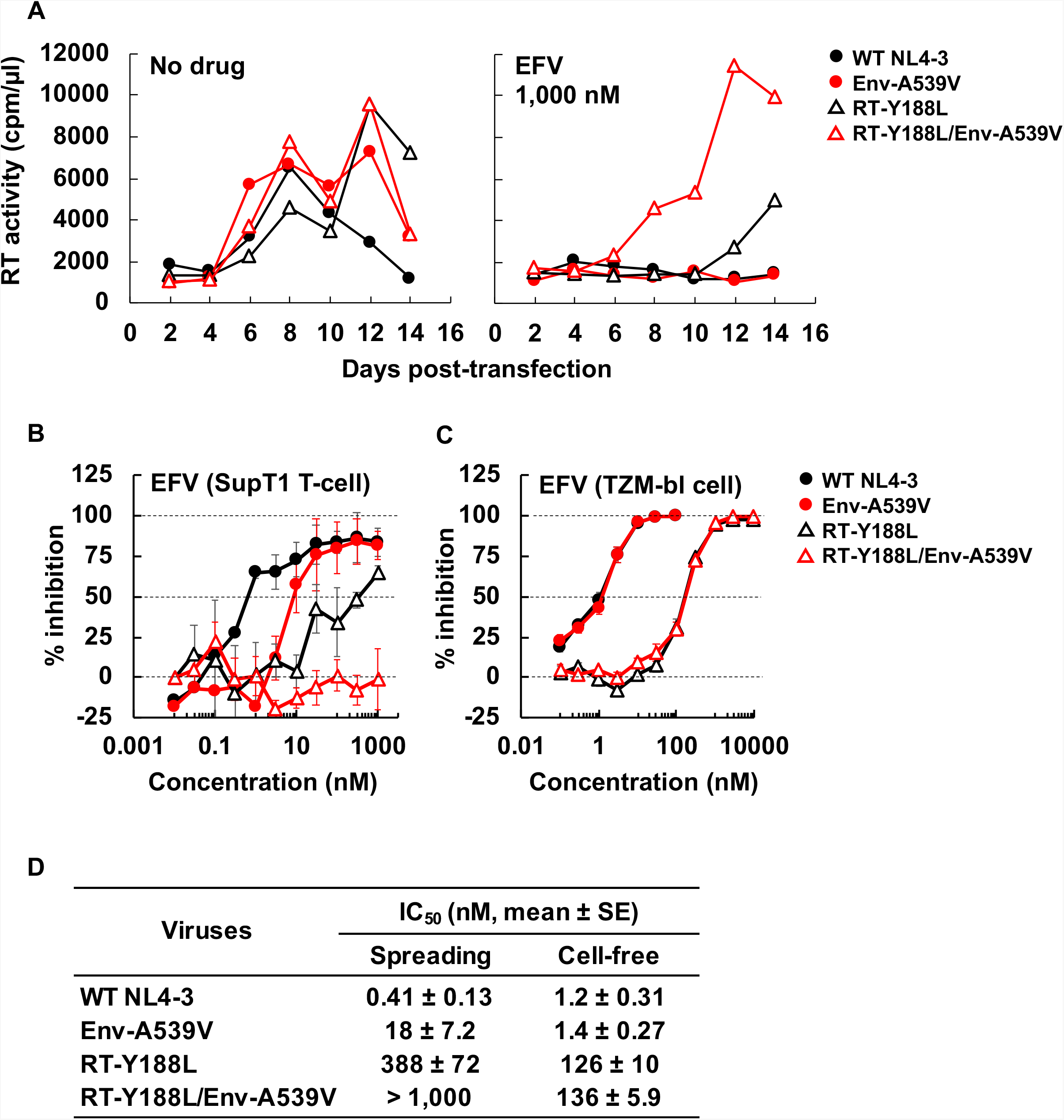
Effect of Env-A539V and RT-Y188L mutations on susceptibility to EFV. (A) The SupT1 T-cell line was transfected with the indicated proviral clones in the absence or presence of 1,000 nM EFV. Virus replication kinetics were monitored by measuring RT activity at the indicated time points. Data are representative of three independent experiments. (B) SupT1 T cells were transfected with the indicated proviral clones in the absence or presence of serial dilutions of EFV (0.03 – 1,000 nM). The dose-dependent inhibition curve was determined based on RT values at the peak of multi-cycle spreading virus replication. (C) TZM-bl cells were exposed to 100 TCID_50_ of the indicated viruses in the absence or presence of serial dilutions of EFV (0.1 – 10,000 nM). Luciferase activity was measured at 48 hrs post-infection. (D) IC_50_ values for EFV in multi-cycle spreading and cell-free infection based on the data in Fig. 4B and C. Data from at least three independent experiments are shown as means ± SE. *p*-values < 0.001 (***), < 0.01 (**), < 0.05 (*) by unpaired *t*-test.

### NL4-3 Env mutations that overcome blocks to HIV-1 replication in spreading infections stabilize the gp120-gp41 interaction

As shown in Fig 1B, our previously reported Env mutations, which we selected for their ability to overcome blocks to virus replication in multi-cycle spreading infections, are located in the C1 domain of gp120 and the HR1 region of gp41. These domains of Env have been shown in mutagenesis and structural studies to be critical for the stability of the gp120-gp41 association in the unliganded state [50–52, 62, 66]. To provide clues regarding the mechanism by which Env mutations enhance the capacity of HIV-1 to spread via cell-cell transfer, we examined their impact on gp120-gp41 association. It is known that sCD4 induces gp120 shedding from virus particles [57]; we therefore incubated purified virions with sCD4 at 37°C for 2 hrs, and then measured the amount of particle-associated gp120 and p24 by western blotting. The amount of gp160 in virions was used as a control, as it is not shed from virions and can be detected with the same antibody used to detect gp120. As expected, levels of gp120 associated with WT NL4-3 particles were decreased in a dose-dependent fashion by sCD4 treatment (0.3 – 10 μg/ml). By contrast, the mutations in both gp120 (Env-Y61H and P81S) and gp41 (Env-A539V and A556T) showed significantly reduced sCD4-induced gp120 shedding (Fig 5A). To quantify this effect, we calculated the 50% effective concentration (EC_50_) of sCD4-induced shedding. While the EC_50_ of sCD4-induced gp120 for WT NL4-3 was 0.99 μg/ml sCD4, the EC_50_ for the Env-P81S and the other Env mutants was 3.5 and > 10 μg/ml, respectively (Fig 5B). To examine the interaction between sCD4 and a representative Env mutant, we measured the sensitivity of Env-A539V infectivity to sCD4 (Fig 5C). Env-A539V showed a 3.7-fold resistance to sCD4 relative to WT. Moreover, the IC_50_ of sCD4 against Env-A539V was 1.1 μg/ml (Table 1), which is more than about 10-fold less than the EC_50_ of sCD4-induced gp120 shedding, suggesting that sCD4 could bind to mutant Env at lower concentrations than needed for gp120 shedding, leading to inhibition of viral entry. To further characterize the intrinsic stability of mutant Envs, we also performed a time-dependent gp120 shedding assay by incubating viruses at 37°C for up to 5 days (Fig 5D). The levels of gp120 on WT NL4-3 particles were decreased during the 5-day incubation period. The Env mutations significantly reduced time-dependent gp120 shedding (Fig 5E). Whereas the half-life of WT NL4-3 gp120 shedding was 2.6 days, that of the Env mutants was > 5 days. These data suggest that the NL4-3 Env mutations stabilize the gp120-gp41 interaction.

**Fig 5.**
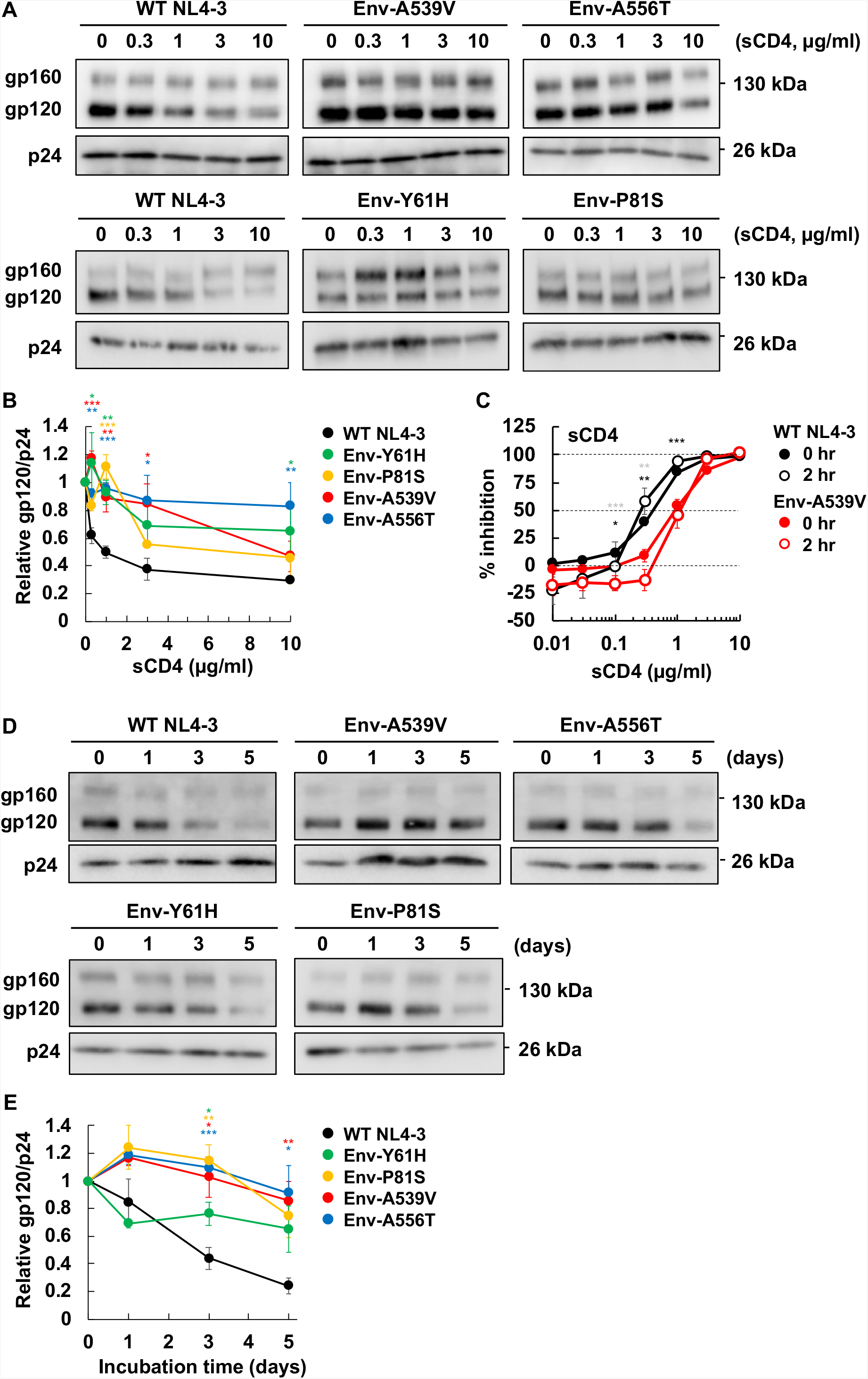
Effect of Env mutations on sCD4-induced and time-dependent gp120 shedding. (A and B) Concentrated viruses were incubated with sCD4 at the indicated concentrations at 37°C for 0 or 2 hrs. Incubated viruses were subsequently purified through a 20% sucrose cushion, and viral proteins were detected by western blotting. A representative gel for the sCD4-induced gp120 shedding assay is shown in A. The ratio of gp120 to p24 was quantified and plotted in B. Data from at least two independent experiments are shown as mean ± SE. (C) TZM-bl cells were exposed to 100 TCID_50_ of the indicated viruses incubated with sCD4 prior to infection for 0 or 2 hrs. Luciferase activity was measured at 48 hrs post-infection. Data from three independent experiments are shown as mean ± SE. *p*-values < 0.001 (***), < 0.01 (**), < 0.05 (*) by unpaired *t*-test, with asterisks for 0 hr or 2 hrs incubation time points indicated in black or gray, respectively. (D and E) Concentrated viruses were incubated at 37°C for the indicated times. Incubated viruses were subsequently purified through a 20% sucrose cushion, and viral proteins were detected by western blotting. A representative gel for the time-dependent gp120 shedding assay is shown in D. The ratio of gp120 to p24 was quantified and plotted in E. Data from at least three independent experiments are shown as mean ± SE. *p*-values < 0.001 (***), < 0.01 (**), < 0.05 (*) by unpaired *t*-test.

### The Env-A539V mutation reduces sensitivity to neutralizing antibodies (NAbs) that recognize the CD4-induced Env conformation

smFRET analysis revealed that the HIV-1 Env trimer samples three conformational states [18, 19]. It has been suggested that anti-Env NAbs and Env-targeting entry inhibitors display preferences for specific Env conformations [18, 19, 67] and that the gp41 ectodomain can modulate the conformational dynamics of Env [52, 62, 66]. To probe the impact of the Env-A539V mutation on Env conformation, we examined the sensitivity of Env-A539V to a panel of NAbs (Fig 6). We used three groups of NAbs: The first group (Figs 6A – E), which included VRC01 (specific for the CD4bs), PGT145, PG16 (specific for the V2 apex), 2G12 and PGT121 (V3-glycan-specific), preferentially bind to the unliganded, closed Env conformation [68–71]. The second group (Fig 6F) included 35O22, which is specific for the gp120-gp41 interface and shows no conformational preference [72]. The third group (Figs 6G - J) included F105 (CD4 bs-specific), 17b (specific for the CD4i epitope), 447-52D (V3 loop), and 10E8 (gp41 MPER), preferentially target the CD4-bound Env conformation [73–76]. Although the Env-A539V mutation is not located within the epitopes of the NAbs used in this analysis, it altered the sensitivity to several NAbs, with reduced sensitivity to F105 (3.1-fold), 447-52D (28-fold), 17b (27-fold), and 10E8 (1.8-fold) and increased (33-fold) sensitivity to PGT145 (Table 1). To further probe the effect of the Env-A539V mutation on Env conformation, we compared NAb binding to Env on 293T cells by flow-cytometry. As indicated in Figs 6M and N, 17b and 10E8 less efficiently bind to Env-A539V relative to WT Env. In contrast, PGT145 and PG16 bind the mutant Env to a similar extent as WT NL4-3 Env (Figs 6O and P). These data suggest that the Env-A539V mutation stabilizes the closed Env conformation. We also examined sensitivity to SIM.4 (an anti-CD4 Ab) [77] and AMD3100 (a CXCR4 antagonist) [78] to evaluate the impact of the Env-A539V mutation on Env function in HIV-1 entry. As shown in Figs 6K and L, Env-A539V showed comparable sensitivity to SIM.4 and AMD3100 compared to WT NL4-3, suggesting that this mutation does not alter CD4 and CXCR4 dependency.

**Fig 6.**
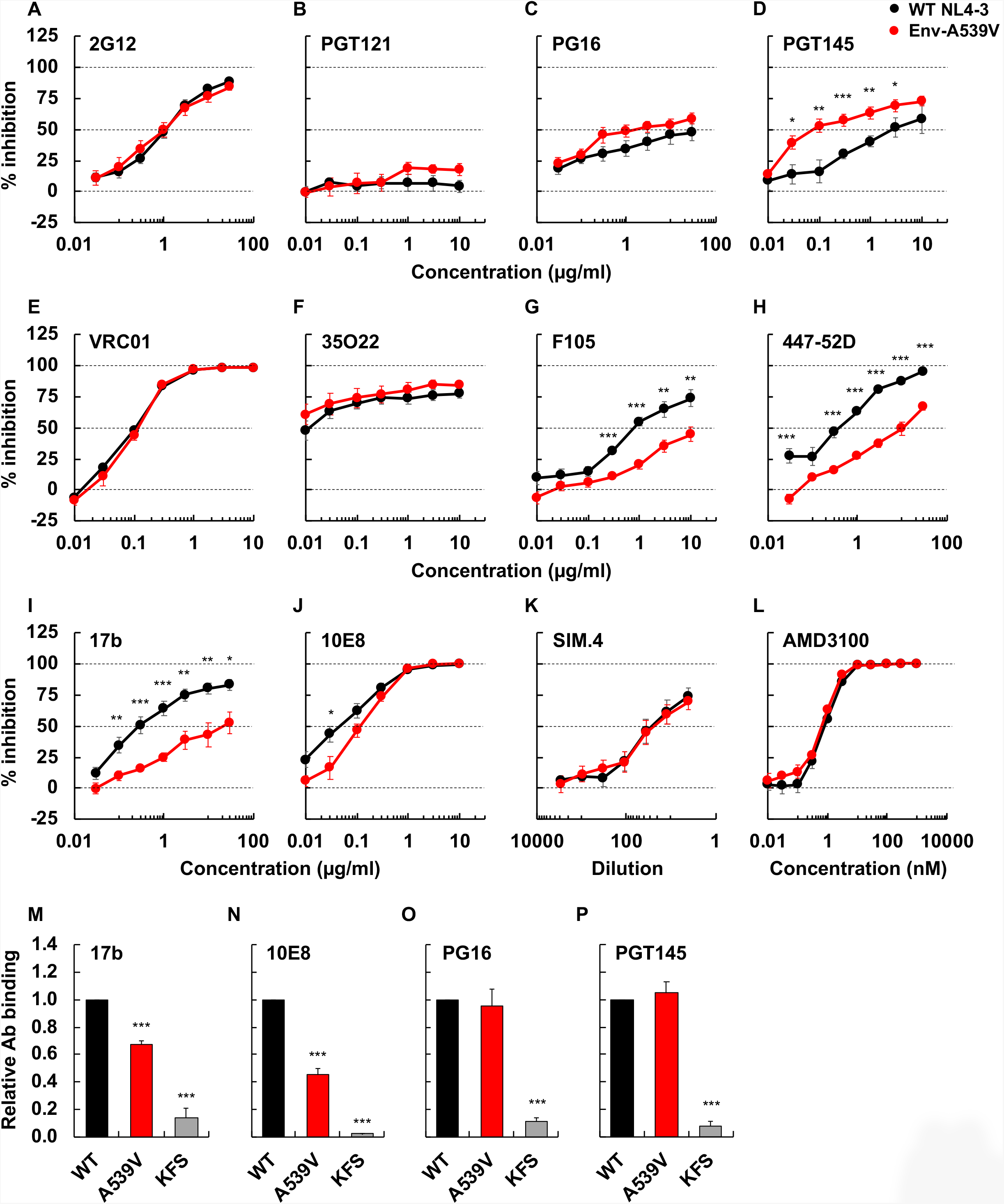
The sensitivity or binding of Env-A539V to NAbs recognizing different Env conformations, anti-CD4 Ab and co-receptor antagonist. TZM-bl cells were exposed to 100 TCID_50_ of WT NL4-3 or Env-A539V viruses in the presence of varying concentrations of NAbs (A - J), anti-CD4 Ab (K) and CXCR4 antagonist (L); luciferase activity was measured at 48 hrs post-infection. Data from three independent experiments are shown as mean ± SE. 293T cells transfected with the GFP-encoding reporter clone pBR43IeG expressing WT NL4-3 Env or the Env-A539V mutant were preincubated with 17b (M), 10E8 (N), PG16 (O) or PGT145 (P) at 4°C for 1 hr. KFS is an Env-defective mutant. The cells were washed, and APC-conjugated anti-human IgG was used to detect bound Ab. APC signals were normalized by GFP signal to calculate the Ab binding efficiency. Data are shown as mean ± SE from three independent experiments. *p*-values < 0.001 (***), < 0.01 (**), < 0.05 (*) by unpaired *t*-test or one-way ANOVA and Tukey’s multiple comparison test.

### Env mutations confer ARV resistance in the context of CCR5-tropic HIV-1

To examine whether Env-mediated resistance to ARVs occurs in strains of HIV-1 other than the CXCR4-tropic, lab-adapted strain NL4-3, we propagated the CCR5-tropic clone NL(AD8) [79] in the SupT1huR5 T-cell line, which expresses high levels of human CCR5, in the presence of DTG. Treatment of transfected SupT1huR5 cultures with 6.0 nM DTG markedly delayed NL(AD8) replication, but at 31 days post-transfection we observed a peak of replication (Fig 7A). Sequencing of the putative escape mutant indicated the absence of any mutations in IN but revealed the presence of an Env-N654K mutation in gp41 HR2 (Fig 7B). Asn at position in 654 is highly conserved (99.76%) in subtype B. To examine whether the Env-N654K mutation in NL(AD8) confers resistance to DTG, we introduced the mutation in WT NL(AD8) and examined replication kinetics in the presence of DTG (Fig 7C). Interestingly, this mutant exhibits faster replication kinetics than WT and can still replicate in the presence of 300 nM DTG (Fig 7C, left panel). IC_50_ calculations indicate that the Env-N654K mutation confers 30-fold resistance to DTG relative to WT NL(AD8) (Fig 7D). As observed with several of the Env mutations selected in our previous study [48], the Env-N654K mutation impairs single-cycle infectivity (Fig 7E) and does not confer DTG resistance in the context of cell-free infection (Fig 7F and Table 2). These results are all consistent with the hypothesis that, as with the mutations selected in the context of NL4-3, the NL(AD8) Env-N654K mutation confers DTG resistance by enhancing the efficiency of cell-cell transfer.

**Fig 7.**
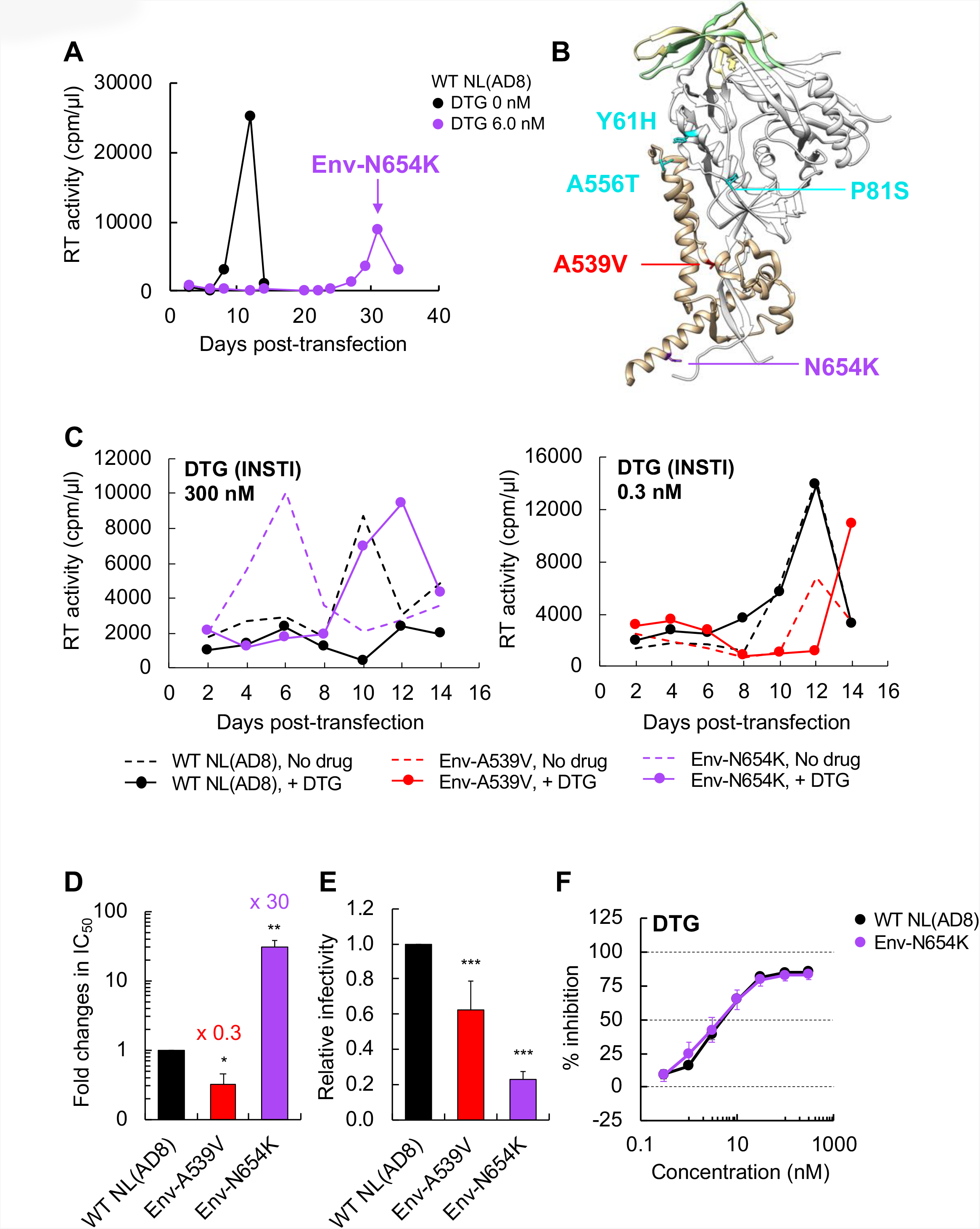
Selection for DTG resistance with the CCR5-tropic NL(AD8) strain. (A) The SupT1huR5 T-cell line was transfected with pNL(AD8) in the absence or presence of 6.0 nM DTG. At the time point indicated by the arrow, DNA was extracted from the DTG-treated culture and the IN- and Env-coding regions were sequenced, leading to the identification of the Env-N654K mutation. (B) A prefusion Env structure of subtype B JR-FL SOSIP.664 (PDB accession number is 5FYK [96]), highlighting the location of Env mutations selected in the context of NL4-3 [48] and the Env-N654K mutation selected in NL(AD8). Env amino acid positions are indicated using the NL4-3 numbering. Most of gp120 is shown in white, with gp120 V1/V2 and V3 loops colored in light yellow and light green, respectively. Env-Y61, P81, and A556 are highlighted in cyan; Env-A539 and N654 are highlighted in red and purple, respectively. The structural model was generated using the UCSF Chimera software. (C) The SupT1huR5 T-cell line was transfected with WT or mutant pNL(AD8) in the absence or presence of 300 or 0.3 nM DTG. Replication kinetics were monitored by measuring RT activity at the indicated time points. Data are representative of at least three independent experiments. (D) The SupT1huR5 T-cell line was transfected with WT or mutant pNL(AD8) in the absence or presence of a serial dilution of DTG (1,000 nM-0.03 nM). DTG IC_50_ values were calculated based on RT values at the peak of virus replication. Fold changes in IC_50_ relative to WT are indicated. Data from at least three independent experiments are shown as means ± SE. *p*-values < 0.001 (***), < 0.01 (**), < 0.05 (*) by unpaired *t*-test. (E) RT-normalized virus stocks produced from HeLa cells were used to infect TZM-bl cells. Luciferase activity was measured at 48 hrs post-infection. (F) TZM-bl cells were exposed to 100 TCID_50_ of the indicated viruses in the presence of various concentrations of DTG and luciferase activity was measured at 48 hrs post-infection. Data are normalized to WT and are shown as means ± SE from at least three independent experiments. *p*-values < 0.001 (***), < 0.01 (**), < 0.05 (*) by unpaired *t*-test.

**Table 2.**
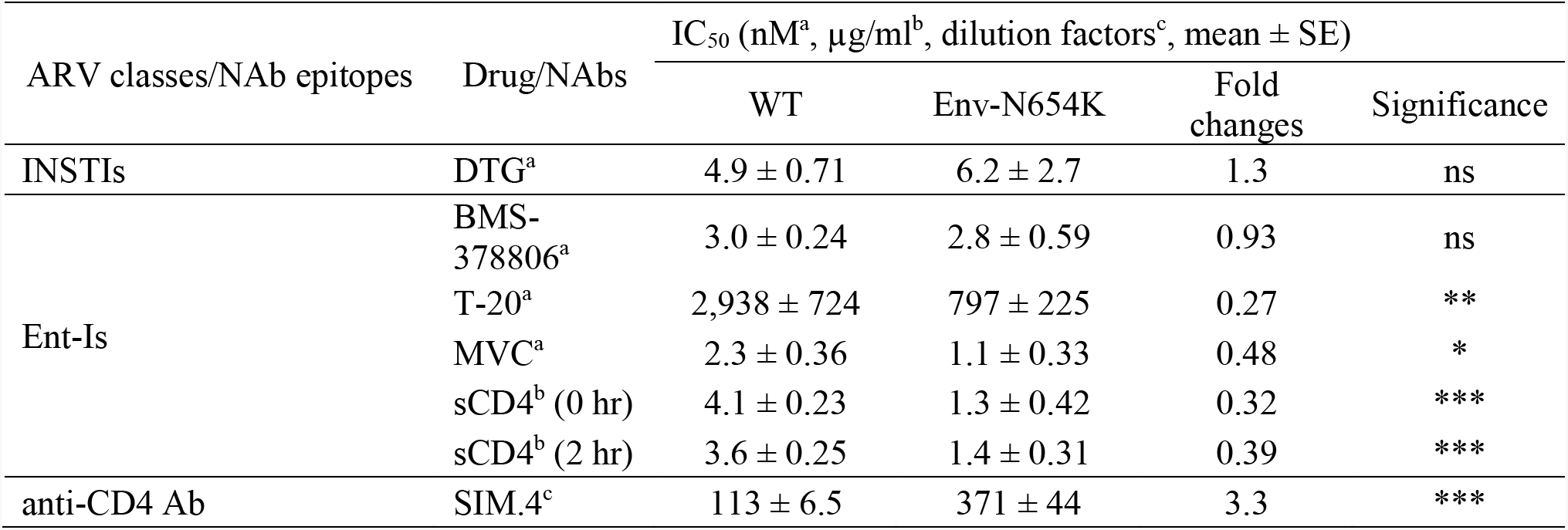
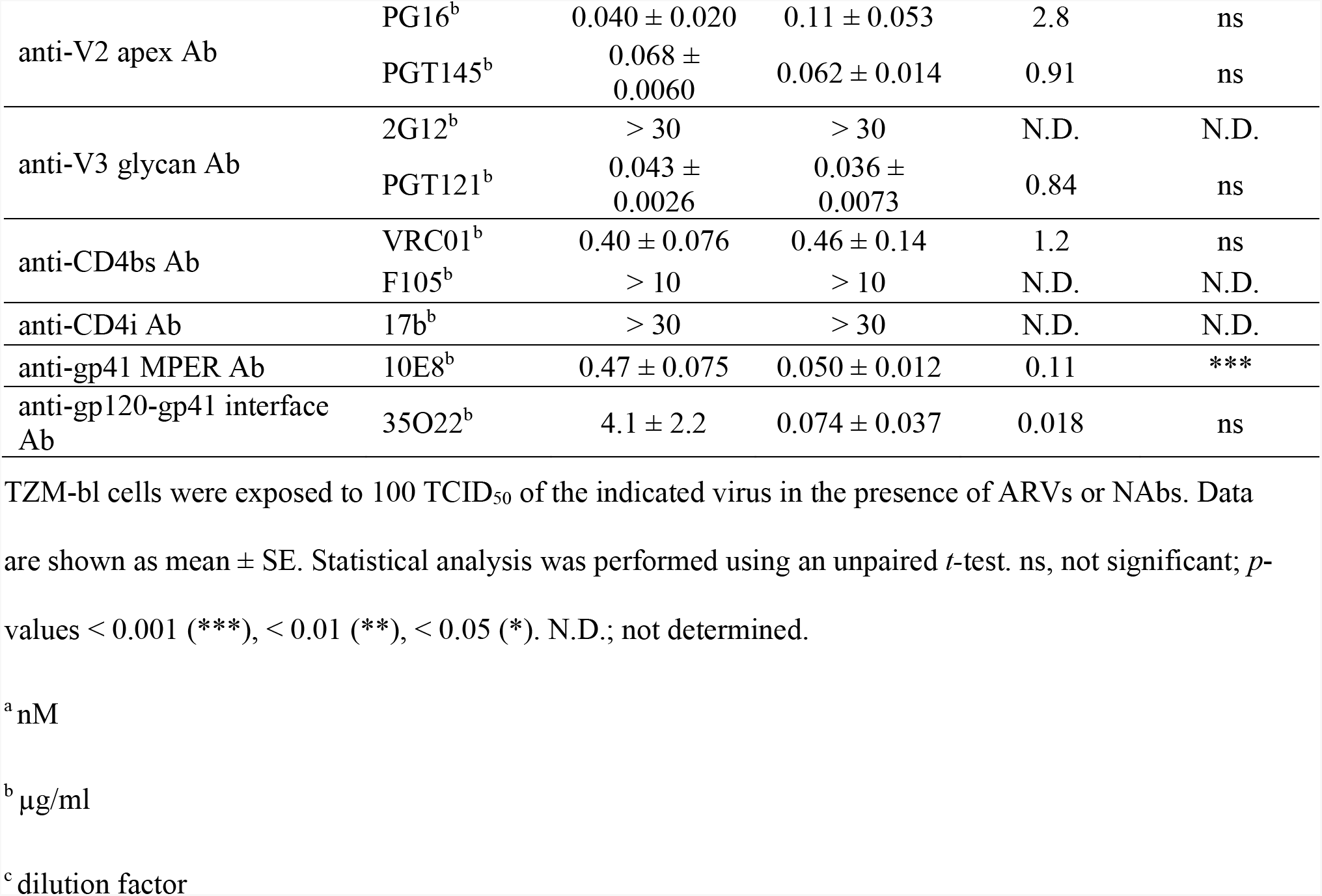
IC_50_ values in cell-free infection for NL(AD8) variants.

| ARV classes/NAb epitopes | Drug/NAbs | IC<sub>50</sub> (nM<sup>a</sup>, µg/ml<sup>b</sup>, dilution factors<sup>c</sup>, mean ± SE) |  |  |  |
| --- | --- | --- | --- | --- | --- |
|  |  | WT | Env-N654K | Fold changes | Significance |
| INSTIs | DTG<sup>a</sup> | 4.9 ± 0.71 | 6.2 ± 2.7 | 1.3 | ns |
| Ent-Is | BMS-378806<sup>a</sup> | 3.0 ± 0.24 | 2.8 ± 0.59 | 0.93 | ns |
|  | T-20<sup>a</sup> | 2,938 ± 724 | 797 ± 225 | 0.27 | ** |
|  | MVC<sup>a</sup> | 2.3 ± 0.36 | 1.1 ± 0.33 | 0.48 | * |
|  | sCD4<sup>b</sup> (0 hr) | 4.1 ± 0.23 | 1.3 ± 0.42 | 0.32 | *** |
|  | sCD4<sup>b</sup> (2 hr) | 3.6 ± 0.25 | 1.4 ± 0.31 | 0.39 | *** |
| anti-CD4 Ab | SIM.4<sup>c</sup> | 113 ± 6.5 | 371 ± 44 | 3.3 | *** |
| anti-V2 apex Ab | PG16<sup>b</sup> | 0.040 ± 0.020 | 0.11 ± 0.053 | 2.8 | ns |
|  | PGT145<sup>b</sup> | 0.068 ± 0.0060 | 0.062 ± 0.014 | 0.91 | ns |
| anti-V3 glycan Ab | 2G12<sup>b</sup> | > 30 | > 30 | N.D. | N.D. |
|  | PGT121<sup>b</sup> | 0.043 ± 0.0026 | 0.036 ± 0.0073 | 0.84 | ns |
| anti-CD4bs Ab | VRC01<sup>b</sup> | 0.40 ± 0.076 | 0.46 ± 0.14 | 1.2 | ns |
|  | F105<sup>b</sup> | > 10 | > 10 | N.D. | N.D. |
| anti-CD4i Ab | 17b<sup>b</sup> | > 30 | > 30 | N.D. | N.D. |
| anti-gp41 MPER Ab | 10E8<sup>b</sup> | 0.47 ± 0.075 | 0.050 ± 0.012 | 0.11 | *** |
| anti-gp120-gp41 interface Ab | 35O22<sup>b</sup> | 4.1 ± 2.2 | 0.074 ± 0.037 | 0.018 | ns |

TZM-bl cells were exposed to 100 TCID<sub>50</sub> of the indicated virus in the presence of ARVs or NAbs. Data are shown as mean ± SE. Statistical analysis was performed using an unpaired *t*-test. ns, not significant; *p*-values < 0.001 (\*\*\*), < 0.01 (\*\*), < 0.05 (\*). N.D.; not determined.
<sup>a</sup> nM
<sup>b</sup> µg/ml
<sup>c</sup> dilution factor

We also propagated the subtype C, CCR5-tropic transmitted-founder isolate K3016 [80] in the presence of DTG. We identified the Env-T529I mutation in gp41 HR1, which is located at the same position as NL4-3 Env-A539V (S1A and B Figs). This mutant showed 2.8-fold-reduced sensitivity to DTG, although its replication was delayed relative to that of WT K3016 (S1C and D Figs). These results indicate that Env-mediated drug resistance *in vitro* can occur in clinically relevant HIV-1 strains independent of co-receptor usage and subtype.

To examine whether the Env-A539V mutation, which was originally selected in the context of NL4-3 [48], would also confer ARV resistance in the context of another viral isolate we introduced this mutation into the NL(AD8) molecular clone. In contrast to the phenotype of this mutant in NL4-3, the NL(AD8) Env-A539V mutant exhibited delayed replication in the absence of DTG and caused an increase (3.1-fold) in DTG sensitivity in multi-cycle spreading infections. Moreover, the cell-free infectivity of the NL(AD8) Env-A539V mutant was reduced relative to that of WT NL(AD8) (Figs 7C - E). These data demonstrate that the impact of Env mutations on replication kinetics and drug sensitivity can be HIV-1 strain-dependent.

Next, to examine the impact of the Env-N654K mutation on the conformation and neutralization properties of NL(AD8) Env, we examined the sensitivity of this mutant to a panel of NAbs (Fig 8 and Table 2). Compared to NL4-3, NL(AD8) is more susceptible to NAbs, such as PGT121, PG16 and PGT145, that preferentially recognize the closed Env conformation (S2 Fig, and Tables 1 and 2). Conversely, NL(AD8) is resistant to 17b (S2 Fig, and Tables 1 and 2) and F105 (Table 1). These results suggest that NL(AD8) tends to sample the closed Env conformation relative to NL4-3. NL(AD8) Env-N654K increased sensitivity to 35O22 (56-fold) and 10E8 (9.1-fold) (Figs 8E and G, and Table 2). Because the effects of N654K on the sensitivity to other NAbs are minimal, this mutation may affect the local structure of HR2 and the MPER rather than the global structural dynamics of NL(AD8) Env. We also examined sensitivity to several entry inhibitors (Figs 8H – L and Table 2). Env-N654K slightly increased sensitivity to SIM.4, maraviroc (MVC; a CCR5 antagonist) and T-20. Moreover, Env-N654K increased the sensitivity to sCD4. Unlike NL4-3 (S2J Fig), incubation time did not affect the susceptibility of Env-N654K to sCD4 (Fig 8H and S2J Fig). This observation suggests that sCD4-induced gp120 shedding is minimal in the NL(AD8) strain, consistent with previous reports for non-lab-adapted isolates [55–57]. Indeed, we did not observe sCD4-induced gp120 shedding from either WT or Env-mutant particles at sCD4 concentrations up to 10 μg/ml (S3A, C and E Figs.). We also examined time-dependent gp120 shedding (S3B and D Figs). In contrast to NL4-3 (Figs 5D and E, and S3F Fig), only minimal gp120 shedding was observed for NL(AD8) Env over a 5-day incubation period at 37°C, and there was no significant difference in shedding between WT and Env-N654K NL(AD8) Env (S3B and D Figs).

**Fig 8.**
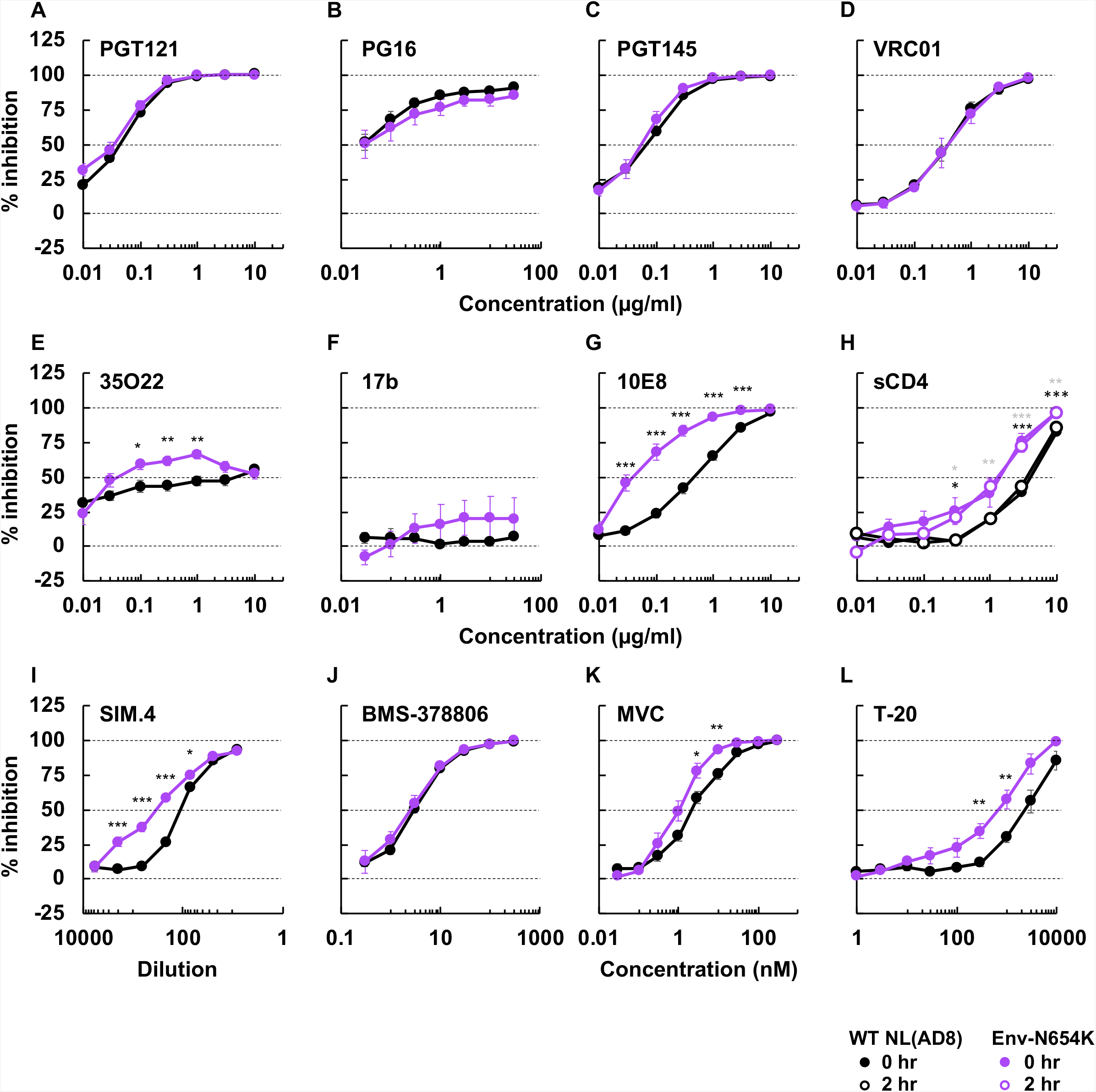
The sensitivity of NL(AD8) Env-N654K to NAbs recognizing different Env conformations, anti-CD4 Ab, and entry inhibitors. TZM-bl cells were infected with 100 TCID_50_ of the indicated viruses in the presence of various concentrations of NAbs (A-G), sCD4 (H), anti-CD4 Ab SIM.4 (I) and entry inhibitors (J – L). Luciferase activity was measured at 48 hrs post-infection. Data from at least three independent experiments are shown as means ± SE. *p*-values < 0.001 (***), < 0.01 (**), < 0.05 (*) by unpaired *t*-test. The asterisks for 0 hr or 2 hrs incubation with sCD4 (panel H) are indicated as black or gray, respectively.

### Sequence analysis of IN- and Env-coding regions of viruses from individuals failing a RAL-containing regimen

Our results indicate that Env-mediated drug resistance can occur in clinically relevant HIV-1 strains *in vitro*. However, whether Env mutations contribute to drug resistance *in vivo* is unclear. To address this question, we used single-genome sequencing to compare viral sequences obtained from the plasma of participants failing a RAL-containing regimen in the SELECT study [5]. The aim of the SELECT study was to examine whether RAL with boosted lopinavir (LPV) would be non-inferior to boosted LPV with NRTIs. By 48 weeks of treatment, 10.3% of 258 participants treated with the RAL-containing regimen experienced virological failure, defined here as the inability to achieve a viral load under 400 copies/ml at two consecutivetime points at or after 24 weeks of treatment. By the end of the follow-up in this study, ten participants developed new resistance mutations in IN, including IN-N155H. In the present study, we compared changes in the sequences of IN- and Env-coding regions from five participants who experienced viral rebound (Table 3). These participants were infected with subtype C, CCR5-tropic strains. Each participant had therapeutic plasma concentrations of RAL (> 33 ng/ml) at the time of virological failure. No resistance mutations in IN were detected at day 0, (Table 3). While we identified the IN-N155H mutation in participant identifier (PID) A5 with high frequency (73%), PID A1, A3 and A5 had resistance mutations in IN with low frequency (Table 3), suggesting that virological failure may have occurred in these participants as a result of mutations elsewhere in the HIV-1 genome.

**Table 3.**
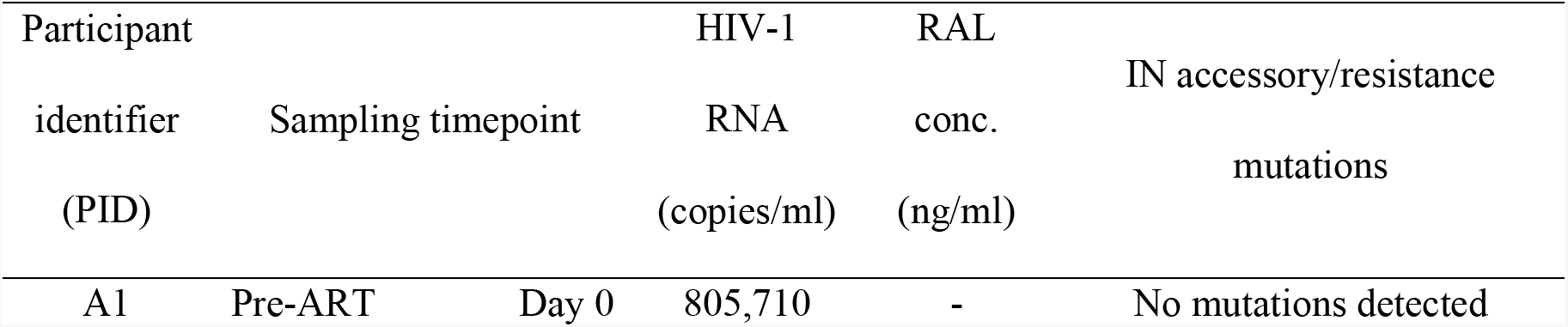
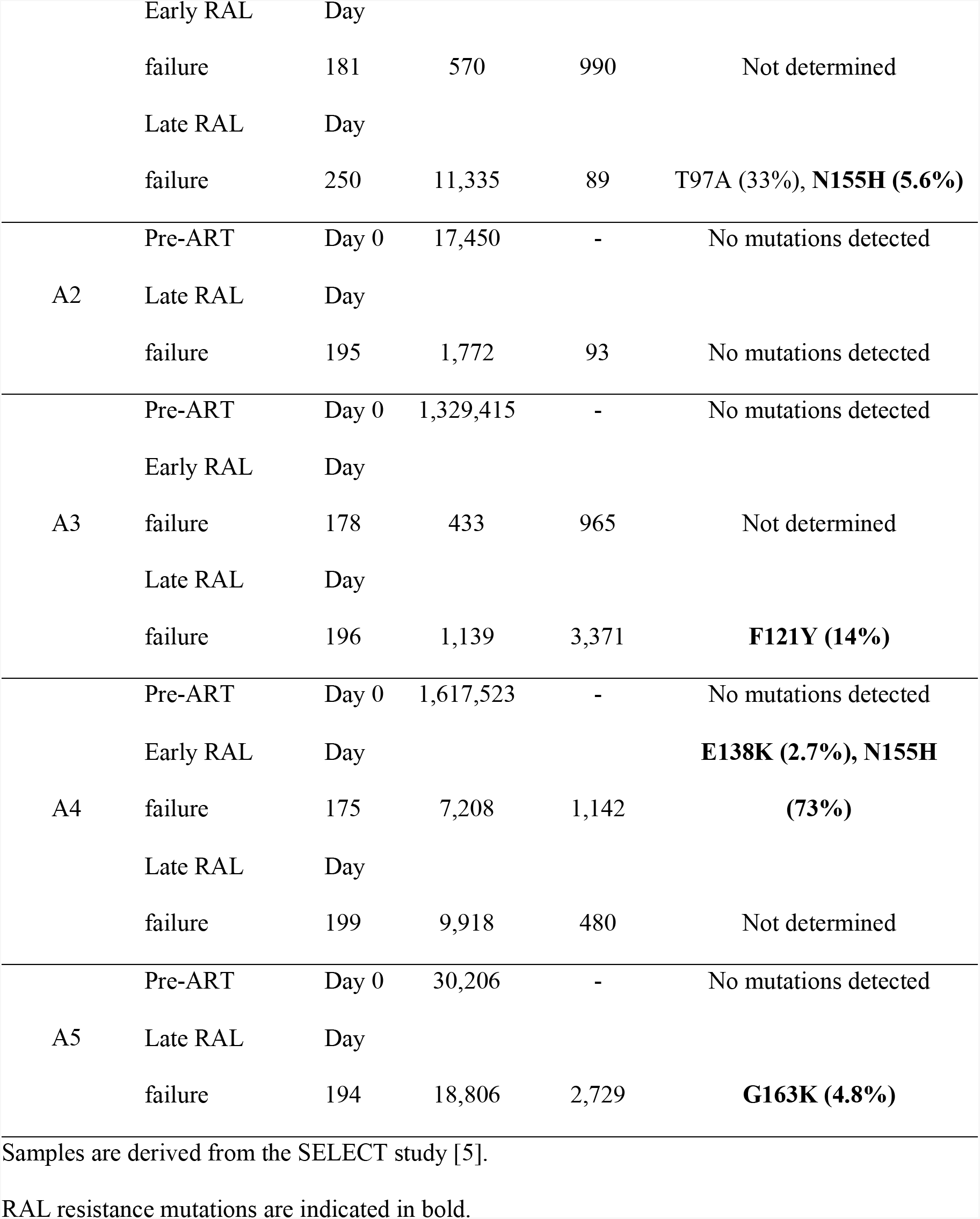
Information on study participants failing a RAL-containing regimen.

Evaluation of the Env-coding region before treatment and during virologic failure indicated the presence of many changes in gp120 and gp41 that emerged or were selected with failure. Most of changes in gp120 were located in highly variable regions, such as the variable loops (V1/V2, V4 and V5 loop), and the C3 region (data not shown). Because these regions contain immunodominant epitopes of subtype C Env, mutations in these regions may have been driven by immune pressure [81, 82]. Although we did not identify mutations at the same positions observed in our *in vitro* studies, we observed an accumulation of mutations in the gp120 C1 domain and gp41 ectodomain in PID A1, A3, A4 and A5 (Figs 9A and B). The frequency of most of the mutations increased significantly with viral rebound. Although we identified a number of Env mutations in PID A2, the frequency of mutations before RAL therapy was similar to that after viral rebound although the RAL-containing regimen decreased viral load before viral rebound (data not shown and Table 3). In many cases, amino acid residues were replaced with more conserved residues; however, we did identify changes to residues with very low prevalence, such as R577K (gp41 HR1) and L602R (gp41 HR2). Some mutated positions in the gp41 ectodomain were identified in several participants (Figs 9A and B). The observation of changes arising in highly conserved positions at the gp120-gp41 interface – a region in which, based on our data, mutations that confer ARV resistance *in vitro* cluster – in the absence of drug-resistance mutations in IN, suggests the possibility that these gp41 changes may have contributed to virological failure in a subset of participants in the study. Further analysis will be needed to explore this hypothesis in more detail.

**Fig 9.**
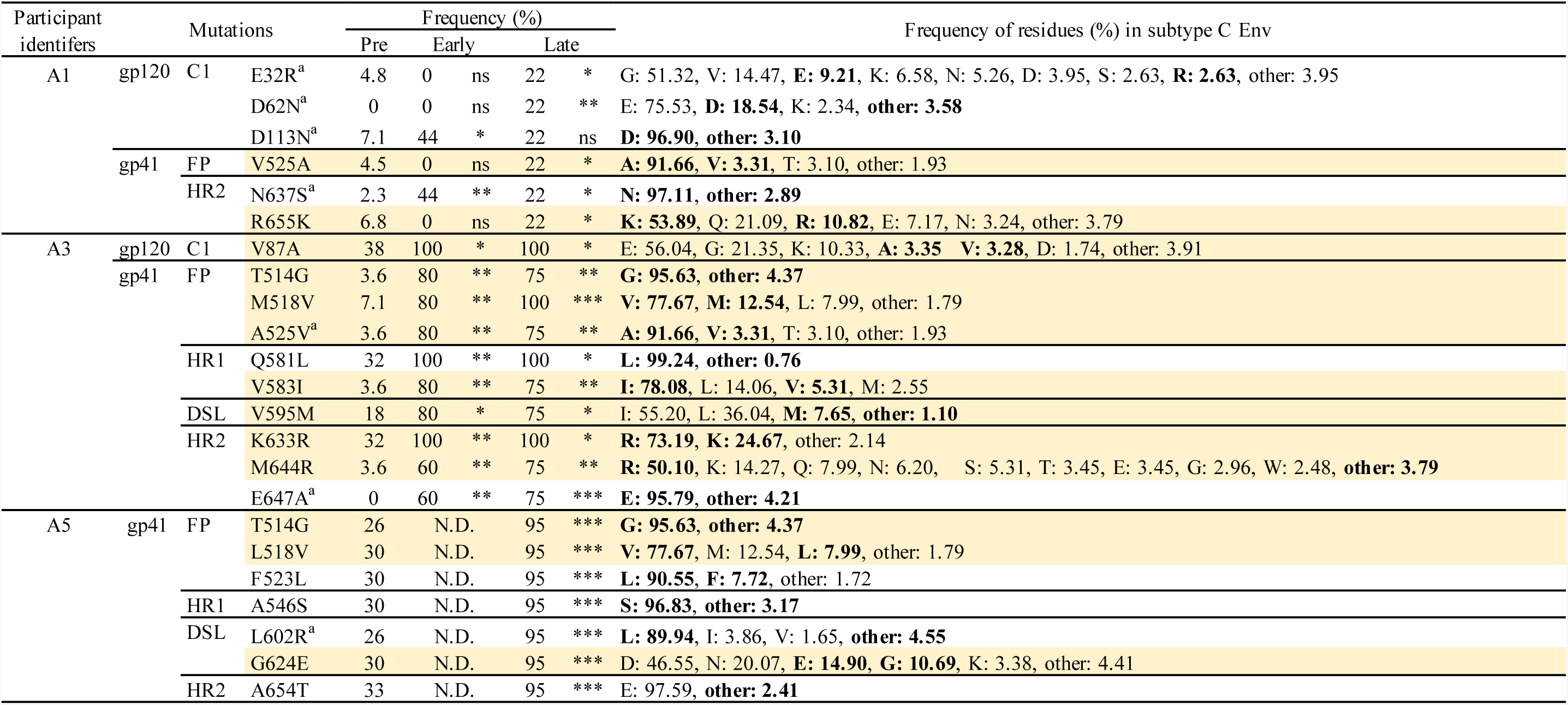
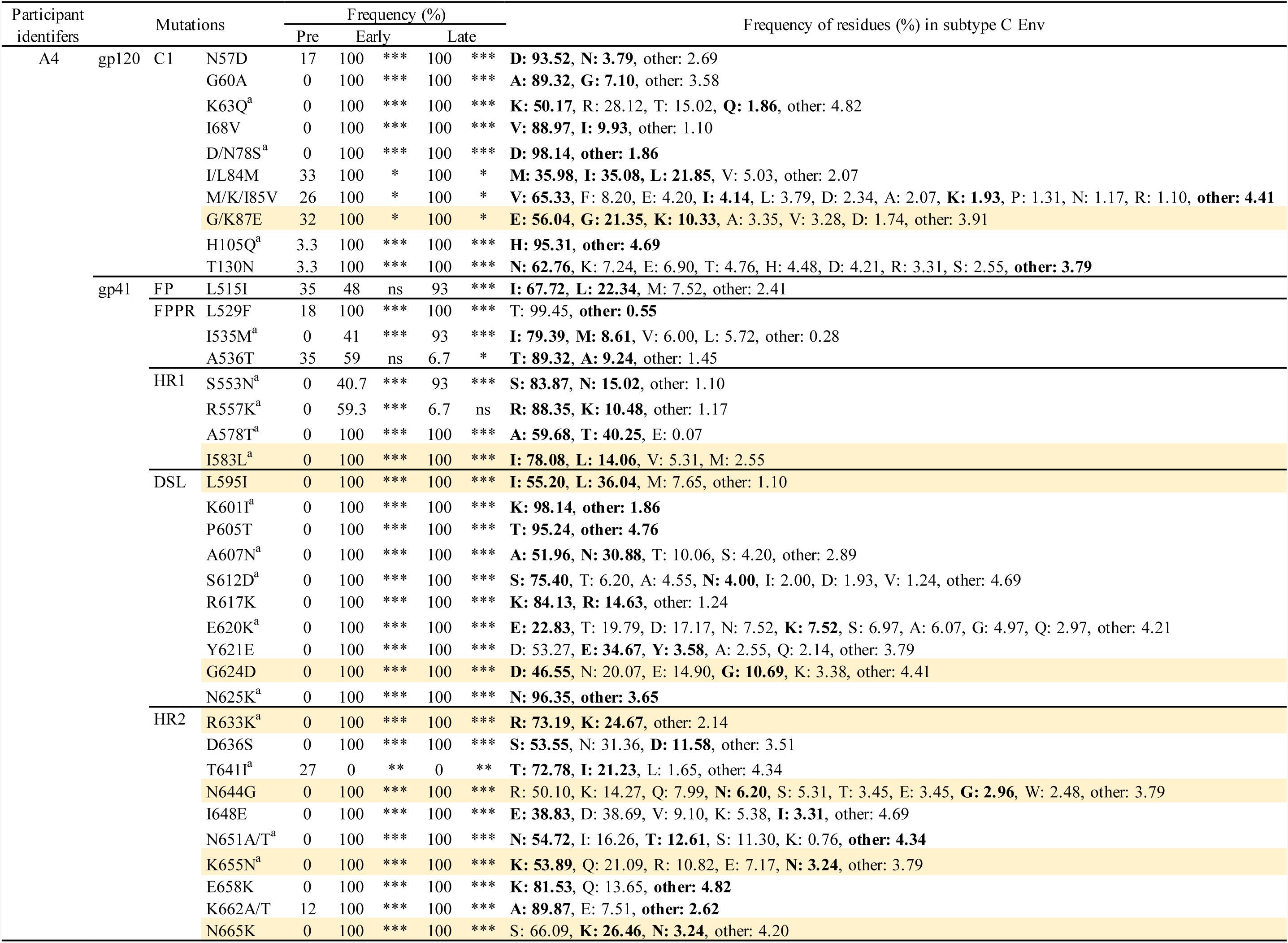
Frequency of observed mutations in gp120 C1 domain and gp41 ectodomain in patient-derived samples from the SELECT study. (A) PID A1, A3 and A5. (B) PID A4. Positions of mutations are indicated using HXB2 numbering. FP, fusion peptide; FPPR, fusion peptide proximal region; HR1/HR2, heptad repeat 1/2; DSL, disulfide loop. Mutations observed in multiple patients are shaded in yellow. Observed residues are indicated in bold. Mutations that changed conserved positions to less conserved residues are indicated with an ^a^. Frequencies were determined for subtype C sequences (n = 5,923) retrieved from the Los Alamos HIV Database. Fischer’s exact test was performed to determine statistical significance. *p*-values < 0.001 (***), < 0.01 (**), < 0.05 (*). ns: not significant, N.D.; not determined.

## Discussion

In this study, we demonstrate that the NL4-3 Env-A539V mutant, which was selected in the presence of DTG, confers broad resistance to multiple classes of ARVs in multi-cycle spreading infections. This Env mutation also increases resistance to ARVs when coupled with ARV target-gene mutations, again in the context of spreading infections. However, the Env mutation does not alter ARV susceptibility in cell-free infection. The results demonstrate that Env mutations clustered at the gp120-gp41 interface can confer broad resistance by enhancing the ability of HIV-1 to spread via a cell-cell route of transmission. Interestingly, NL4-3 Env mutants exhibiting an enhanced ability to spread via cell-cell transmission have more stable and closed Env conformations compared to WT NL4-3. By propagating CCR5-tropic HIV-1 strains NL(AD8) and K3016 in the presence of DTG, we obtained gp41 ectodomain mutations Env-N654K and Env-T529I, respectively. We demonstrate that these Env mutations reduce susceptibility to DTG in multi-cycle spreading infection as observed for the NL4-3 Env mutants, indicating that Env-mediated drug resistance can occur in non-laboratory-adapted HIV-1 strains. However, the effects of the Env mutations on replication kinetics and Env structure are HIV-1 strain-dependent. Finally, we performed single-genome sequencing analysis of the IN- and Env-coding regions of viruses from infected individuals failing a RAL-containing regimen. We observed that many mutations accumulated in the Env- but not IN-coding region in most study participants. While most of the gp120 changes that arose *in vivo* were located in highly variable regions, a number of changes in the gp41 ectodomain were observed at the highly conserved gp120-gp41 interface, as observed *in vitro*.

Several studies have demonstrated that cell-cell transfer is associated with a reduced susceptibility to ARVs relative to cell-free infection [32–34]. We previously proposed a model, consistent with other findings [33], whereby Env mutations that enhance cell-cell transmission increase the MOI following viral transfer across the VS; concentrations of DTG that are sufficient to inhibit cell-free infection by these mutants are therefore insufficient to block their cell-cell transmission [48]. Based on this model, the selected Env mutations would be predicted to confer resistance to multiple ARVs independent of their mode of action, and should confer resistance in the context of a spreading, but not cell-free, infection. Indeed, the NL4-3 Env-A539V mutant selected in the presence of DTG exhibits broad resistance to a number ARVs including INSTIs, NRTIs, NNRTIs and PIs in spreading replication assays but not in cell-free infectivity assays. These observations suggest that the broad resistance conferred by the HIV-1 Env mutations will affect susceptibility to not only the currently approved ARVs, but also potentially to next-generation drugs.

In contrast to the other classes of ARVs, Env-A539V is sensitive to two types of Ent-Is: a fusion inhibitor (T-20) and an attachment inhibitor (BMS-378806). Previous studies have shown that Ent-Is targeting CD4 or co-receptor are effective in blocking cell-cell transmission [34, 60, 83]. While some CD4bs-targeting agents inefficiently inhibit cell-cell transmission, anti-gp41 MPER Abs and T-20 are effective [60, 84]. These observations suggest that accessibility of epitopes or time of action may be important factors for the efficacy of Ent-Is in blocking cell-cell transfer. Interestingly, the Env-A539V mutation has been reported to emerge in the presence of low concentrations of fusion inhibitors *in vitro*; however, this mutation does not confer resistance to the fusion inhibitor and reverted back to WT at high inhibitor concentration [85]. Overall, our findings imply that including Env-targeted inhibitors in a cART regimen may help prevent the emergence of the type of broadly resistance-conferring Env mutations described here and previously [48].

Virological failure has been observed in patients on PI-containing regimens in the absence of PI-resistance mutations in PR or in the vicinity of Gag cleavage sites [7, 86–89]. Likewise, recent studies have demonstrated that viral rebound can occur in individuals on INSTI-containing therapies without the emergence of INSTI-resistance mutations in IN [5, 6]. In some studies [90, 91] there are concerns about whether the infected individuals remained compliant (i.e., whether suppressive concentrations of the inhibitors were maintained), but in other studies (e.g., La Rosa et al., 2016 [5]) the maintenance of adequate plasma drug concentrations was experimentally verified. In some cases in which target-gene mutations were identified in the treated individuals experiencing virological failure, the resistance mutations identified conferred only low-level resistance [3–10]. These findings suggest that mutations outside the *pol*-coding region (and not around Gag cleavage sites, in the case of PIs) also contribute to virological failure in association with the target-gene mutations. Mutations in the cytoplasmic tail of gp41 have been previously proposed to contribute to PI resistance *in vivo* [13], potentially reflecting the role of virion maturation in Env-mediated viral entry [92, 93], and mutations in the gp41 HR region have also been observed in association with PI failure although the contribution of the mutants to virological failure is still unclear [9]. In addition to mutations in IN, it has also been reported that mutations in the 3’ PPT contribute to INSTI failure both *in vitro* and *in vivo* [12, 94]. We demonstrate here that the Env-A539V mutation rescues the defect in virus replication imposed by the RT-Y188L mutation [65] and increases the resistance of this RT mutant to EFV in multi-cycle spreading but not cell-free infection. These observations indicate that Env mutations can increase resistance conferred by mutations in the ARV-target gene by enhancing viral cell-cell transmission. Mathematical modeling suggests that cell-cell transmission increases the probability that NAb-resistance mutations will emerge, relative to cell-free infection, as a result of reduced NAb potency in the context of cell-cell transmission [84]. As shown in Fig. 2, Env-A539V provides a replication advantage over WT NL4-3 in the presence of ARVs, suggesting that mutations in Env such as those described here may facilitate the acquisition by the virus of resistance mutations in ARV-target genes. According to this “stepping stone” model, by facilitating virus replication in the presence of ARVs the Env mutations would facilitate the emergence of high-level resistance mutations in ARV-target genes. Additional studies will be needed to investigate the role of HIV-1 Env in the development of high-level drug resistance.

We observed that Env mutations in the NL4-3 strain reduce sCD4-induced and time-dependent shedding of gp120 from viral particles. In addition, the Env-A539V mutation decreases sensitivity to NAbs preferring the CD4-bound conformation, whereas this mutant is more susceptible to PGT145, which recognizes the closed Env conformation. These observations suggest that the NL4-3 mutant Envs are more closed and stable on viral particles compared to WT NL4-3 Env. Flow-cytometry analysis using Env-expressing cells indicates that the NAb-binding properties of the mutants largely parallel their neutralization properties, suggesting that mutant Envs on the surface of infected cells exhibit similar conformational dynamics to those on viral particles. Because the NL4-3 Env-A539V mutation does not alter sensitivity to an anti-CD4 Ab or a CXCR4 antagonist, it is likely that the enhancement of cell-cell transfer is due to the increased probability of gp120-CD4 interactions, rather than the increased affinity of gp120 for CD4 or CXCR4. Our findings suggest that decreased gp120 shedding may contribute to enhanced Env-CD4 interactions at cell-cell contact sites. To better understand how Env mutations stabilize gp120-gp41 interactions, we generated structural models using the published Env structures (S4 and S5 Figs) [95, 96]. The models suggest that, relative to the WT, Env-A539V and A556T likely make additional gp120-gp41 contacts in the unliganded Env structure. In addition, the His at residue 61 in the Env-Y61H mutant is predicted to form a salt bridge with E558 in the gp41 HR1 domain (S4 Fig). These new contacts may contribute to stabilization of the gp120-gp41 interaction. In contrast, the Env-P81S mutation does not result in the formation of obvious new contacts in either the unliganded or CD4-bound conformation; however, this mutation disrupts the interaction between Env-P81 and Env-P79 in the CD4-bound conformation (S5 Fig). These two residues are located in a loop region of the α0 helix in the CD4-bound conformation. This helix is formed after CD4 binding and is suggested to have important roles in conformational rearrangements of Env [95]. Based on the results of our gp120 shedding assays, we hypothesize that the Env-P81S mutation may affect the folding of the CD4-bound gp120 structure, resulting in the mutant Env adopting a more closed conformation and reducing gp120 shedding. Our results provide new insights into the relationship between cell-cell transfer and the stability of the gp120-gp41 interaction.

We performed resistance analyses using the more clinically relevant HIV-1 strains NL(AD8) and K3016. We introduced the Env-A539V mutation into NL(AD8) and observed a different phenotype from that observed with this mutation in NL4-3. Whereas NL4-3 Env-A539V exhibited a faster-than-WT replication kinetics, NL(AD8) Env-A539V exhibited impaired replication kinetics and was more susceptible to DTG than WT NL(AD8). These results imply that the Env-A539V mutation will not emerge in the context of the NL(AD8) strain. However, DTG selection experiments led to the emergence of an Env-T529I mutation, which is located at the same position as NL4-3 Env-A539V, in the context of the subtype C, CCR5-tropic transmitted-founder isolate K3016. Although this mutant replicated more slowly than WT K3016 it exhibited reduced susceptibility to DTG. These results highlight the context dependence of resistance-conferring Env mutations, consistent with the suggestion that residues in the gp41 ectodomain are functionally linked with gp120 and modulate conformational dynamics of Env in a strain-dependent manner [52, 62, 97, 98]. As shown in S2 and S3 Figs, the stability of the gp120-gp41 interaction and the conformational dynamics of NL(AD8) Env are different from those of NL4-3 Env, suggesting that the differences in intra- and interprotomer interactions of Env are likely to affect the viral phenotypes of the Env mutants in terms of their fitness and ability to confer broad ARV resistance.

Propagation of NL(AD8) in the presence of DTG led to the selection of the Env-N654K mutation in gp41 HR2. Like several of the selected NL4-3 Env mutations (this report and Van Duyne et al., 2019 [48]), the NL(AD8) Env-N654K mutation confers resistance to DTG and is highly fit in multi-cycle spreading infections, but is also profoundly defective in cell-free infectivity. These observations suggest that this mutant spreads predominantly via a cell-cell route as observed for the NL4-3 Env mutants. However, the effects of the NL(AD8) Env-N654K mutation on Env stability and conformational dynamics appear to be different from those of the NL4-3 Env mutants. This mutation increases sensitivity to the anti-CD4 Ab, MVC, and to sCD4, T-20 and anti-MPER Ab, suggesting that this mutation may alter Env conformational changes after CD4 binding. An Asn at Env-654 is highly conserved (99.76% in subtype B), and previous studies have shown that substitutions at this position decrease the affinity of HR2 for HR1, resulting in decreased cell-free infectivity and fusogenicity [99–101]. Several lines of evidence suggest that syncytium formation is tightly regulated at the site of cell-cell contact by viral and cellular factors [92, 93, 102]. Although the roles of syncytium formation in HIV-1 replication are still debated, the formation of large syncytia may negatively impact HIV-1 replication [102, 103]. Several studies have reported that mutations in Env arose in infected SupT1 cells and improved replication in the absence of syncytium formation [104–107]. As we previously reported, Env-Y61H, P81S and A556T in the NL4-3 strain enhance cell-cell transmission while impairing cell-free infectivity and fusogenicity, indicating that for these mutants reduced fusogenicity is associated with enhanced cell-cell transmission capacity [48]. Additional studies will be needed to elucidate the precise mechanism of action of the Env-N654K mutation.

To explore the possibility that Env-mediated ARV resistance may occur *in vivo*, we performed single-genome sequence analysis using samples from the SELECT study [5], in which virological failure occurred despite the maintenance of therapeutic levels of RAL in the plasma. Because INSTIs are highly potent ARVs, several groups have investigated whether INSTI monotherapy would effectively suppress viral loads in animal models and human participants [6, 108, 109]. These trials demonstrated that INSTI monotherapy is not sufficient to suppress viral replication and viral loads rebounded. However, in many cases, virological failure occurred in either the absence of mutations in IN, or in association with low-frequency resistance mutations in IN [110]. Consistent with these observations, our single-genome sequence analysis revealed that four of the five participants we analyzed had only low-frequency resistance mutations in the IN-coding region. These results suggest that virological failure occurred in these four participants as a result of mutations outside the Pol-coding region. As expected, we observed that many Env mutations accumulated in less-conserved regions of Env, such as gp120 C3 and variable loops, which are predominant targets for the Ab response [81, 82]. More interestingly, we also found that several mutations in the gp41 ectodomain accumulated at the highly conserved gp120-gp41 interface, as observed in our *in vitro* selection experiments, despite the fact that none of these changes was identical to those selected *in vitro*. As shown in our *in vitro* studies, it is unlikely that the identical mutations observed *in vitro* would emerge *in vivo* because the effects of the Env mutations on viral fitness and drug resistance is viral strain dependent. It is interesting to note that viruses from PID A4, which had the IN-N155H mutation with high frequency, had many changes in the gp120 C1 domain and the gp41 ectodomain compared to viruses isolated from the other participants. A limitation of the current study is the relatively small number of INSTI-failure samples analyzed. Larger numbers of samples will be needed to identify specific mutational patterns associated with virological failure. In addition, other regions outside IN, such as the 3’ PPT, may also be associated with virological failure [12, 94]. Nevertheless, our findings provide tantalizing support for the possibility that HIV-1 Env simultaneously evolves to escape from both NAbs and ARVs in association with ARV target-gene mutations *in vivo*.

In summary, the findings of this study demonstrate that mutations in Env can contribute to broad HIV-1 drug resistance *in vitro*. These findings offer mechanistic insights into Env-mediated drug resistance *in vitro* and provide new clues to understand how HIV-1 develops high-level resistance to ARVs *in vivo*.

## Materials and Methods

### Cell lines

HeLa, HEK293T, and TZM-bl cells [111] were maintained in Dulbecco’s modified Eagle medium (DMEM) supplemented with 10% fetal bovine serum (FBS) at 37°C in 5% CO_2_. The SupT1 and SupT1huR5 T-cell lines [112] were cultured in RPMI-1640 medium supplemented with 10% FBS at 37°C in 5% CO_2_. The SupT1huR5 T-cell line was a kind gift from James Hoxie, Perelman School of Medicine, University of Pennsylvania, Philadelphia, PA.

### Compounds and neutralizing antibodies

DTG and BMS-378806 were purchased from MedChemExpress. RAL, FTC, EFV, NFV, T-20, AMD3100 [78], MVC, SIM.4 [77], sCD4, PGT145 [69], PG16 [70], 2G12 [68], PGT121 [69], VRC01 [71], 447-52D [74], 17b [76], 10E8 [75], F105 [73], 35O22 [72], 16H3 [113], and HIV-Ig were obtained from the NIH AIDS Reagent Program, Division of AIDS, NIAID, NIH. BI-224436 [58] was a kind gift from Alan Engelman, Dana Farber Cancer Institute, Boston, MA.

### Cloning and plasmids

The full-length HIV-1 molecular clones pNL4-3 [114] and pNL(AD8) [79], and the subtype C transmitted founder viral clone CH185 (here denoted K3016, a kind gift from Christina Ochsenbauer and John Kappes, University of Alabama) [80] were used in this study. pBR-NL43-IRES-eGFP-nef^+^ (pBR43IeG), a proviral clone that coexpresses Nef and eGFP from a single bicistronic RNA, was obtained from Frank Kirchhoff through the NIH AIDS Reagent Program [115, 116]. pNL4-3 and pBR43IeG clones bearing Env mutations (Env-Y61H, P81S, A539V and A556T) were described previously [48]. The pBR43IeG/KFS clone, which does not express HIV-1 Env, was described previously [48, 117]. pNL4-3 RT-Y188L [65] was constructed by an overlap PCR method using *SpeI* and *AgeI* restriction sites. pNL(AD8) Env mutants Env-A539V and N654K were constructed by an overlap PCR method using *BamH*I and *EcoR*I restriction sites. Constructed plasmids were verified by Sanger DNA sequencing (Psomagen).

### Preparation of virus stocks

The HEK293T and HeLa cell lines were transfected with HIV-1 proviral DNA using Lipofectamine 2000 (Invitrogen). At 48 hrs post-transfection, virus-containing supernatants were filtered through a 0.45 μm membrane (Merck Millipore). The amount of virus in the supernatants was quantified by RT assay. RT assays were performed as described previously [118] with minor modification. Briefly, after incubation of the virus supernatants with RT reaction mixtures, which contained a template primer of poly (rA) (5 μg/ml) and oligo-dT12-18 primers (1.57 μg/ml) in 50 mM Tris, pH 7.8, 75 mM KCl, 2 mM dithiothreitol, 5 mM MgCl_2_, 0.05% Nonidet P-40, and 0.25 μCi of ^32^P-dTTP at 37°C for 3 hrs, the mixtures were spotted onto filtermat B (Perkin Elmer, cat#1450-521) soaked in 0.5% (v/v) branched-polyethylenimine (Merck Millipore, cat#402727). After washing the spotted filtermat with 2 x SSC buffer (300 mM NaCl and 30 mM sodium citrate), levels of bound ^32^P were measured on a Wallac BetaMax plate reader (Perkin Elmer). TCID_50_ of the virus stocks was determined using TZM-bl cells.

### Virus replication kinetics assays

Virus replication was monitored in SupT1 cells as previously described with minor modifications [48]. Briefly, SupT1 cells were incubated with pNL4-3 clones (1.0 μg/1.0 x 10^6^ cells) in the presence of 700 μg/ml DEAE-dextran at 37°C for 15 min. Transfected cells (1.5 x 10^5^ cells) were plated in 96 well flat-bottom plates and incubated at 37°C in the presence of various concentrations of ARVs. For CCR5-tropic viruses, we used SupT1huR5 cells, which express high levels of human CCR5. Aliquots of supernatants were collected to monitor RT activity, and cells were split 1:3 every other day with fresh drug and media. IC_50_ values were calculated based on RT activity at the peak of replication of each virus. To identify the selected mutations in Pol/Env-coding region, genomic DNA was extracted from infected cells using DNeasy Blood & Tissue mini kit (Qiagen), and then the Pol and Env-coding regions were amplified by PrimeSTAR GXL DNA Polymerase (Takara) and sequenced (Psomagen) using previously described primers [48].

### Single-round infection assays

For cell-free infectivity assays, TZM-bl cells (1.0 x 10^4^ cells) in 96-well plates were exposed to RT-normalized virus stocks produced in HeLa cells. For drug sensitivity assays, 100 TCID_50_ of virus produced in 293T cells was exposed to TZM-bl cells (1.0 x 10^4^ cells) in the presence of various concentrations of drugs or neutralizing antibodies. For sCD4 sensitivity assay, prior to the infection, the viruses were incubated with various concentration of sCD4 at 37°C for 0 or 2 hrs. For assays using PIs or ALLINIs, viruses were produced in the presence of serial dilutions of inhibitors as described previously [119]. Briefly, 293T cells were transfected with WT or mutant molecular clones as indicated above. 6 hrs post-transfection, 1.5 x 10^5^ transfected cells were incubated with a 3-fold serial dilution of the inhibitors at 37°C for 48 hrs. Virus-containing supernatants were harvested and used to infect TZM-bl cells. At 48 hrs post-infection, luciferase activity was measured using the Britelite plus reporter gene assay system and Wallac BetaMax plate reader (Perkin Elmer).

### gp120 shedding assay

To analyze time-dependent gp120 shedding, viruses produced from HeLa cells were incubated at 37°C for 0, 1, 3, or 5 days. For the sCD4-induced gp120 shedding assay, concentrated viruses were incubated with sCD4 (0, 0.3, 1.0. 3.0, and 10 μg/ml) at 37°C for 2 hrs. Following incubation, the viruses were purified by ultracentrifugation through 20% sucrose cushions (60,000 x g) for 45 min at 4°C. The levels of virion-associated gp160, gp120, and p24 were determined by western blotting.

### Western blotting

Viral proteins were separated by SDS-PAGE and transferred to polyvinylidene disulfide (PVDF) membranes (Merck Millipore). After blocking the membranes with 5% skim milk, they were probed with primary antibodies for 1 hr, and then incubated for 1 hr with species-specific horseradish peroxidase-conjugated secondary antibody. After the final washes, bands were detected by chemiluminescence with a Sapphire Biomolecular Imager (Azure Biosystems). Quantification was performed using ImageStudio Lite (Li-Cor Biosciences) software. The p24 protein was detected with anti-HIV-Ig at a final concentration of 5.0 μg/ml in 5% skim milk. Gp160 and gp120 were detected with the 16H3 monoclonal antibody at a final concentration of 0.5 μg/ml in 5% skim milk.

### Antibody binding assay

HEK293T cells were transfected with pBR43IeG constructs as indicated above. At 24 hrs post-transfection, the cells were detached with 5 mM EDTA/PBS, and then washed with PBS. 2.0 x 10^5^ cells were incubated with the specified antibodies at a final concentration of 1.0 μg/ml at 4°C. After 1 hr incubation, the cells were washed with PBS. The cells were then washed with PBS and incubated with allophycocyanin (APC)-conjugated F(ab′)2 fragment donkey anti-human IgG antibody (Jackson ImmunoResearch Laboratories, cat# 709-136-149) at a final concentration of 62.5 μg/ml. The cells were washed with PBS and fixed with 4% paraformaldehyde (PFA, Boston Biosciences). Fixed cells were analyzed with BD LSR-II (BD Biosciences). Data were analyzed by FCS Express Cytometry Software 7 (De novo Software).

### Ethics Statement

Plasma samples were obtained prior to and after virologic failure from donors on combination ART of ritonavir (RTV)-boosted lopinavir (100 mg RTV, 400 mg LPV) plus 400 mg RAL twice a day. Samples were collected by the AIDS Clinical Trial Group (ACTG) study A5273 - a randomized, open-label, phase 3, non-inferiority study at 15 research sites in nine resource-limited countries (three sites in India and South Africa, two in Malawi and Peru, and one each in Brazil, Kenya, Tanzania, Thailand, and Zimbabwe) [5]. The primary endpoint was time to confirmed virologic failure (two measurements of HIV-1 RNA viral load >400 copies/ml). This trial was registered with ClinicalTrials.gov, NCT01352715. Entry and exclusion criteria are listed in the protocol [5]. The study was approved by ethics committees at each site and written informed consent was obtained from each participant.

### Plasma viral RNA extraction and cDNA synthesis

Viral RNA was extracted from plasma containing 200-20,000 copies of HIV-1 RNA. Plasma was first centrifuged at 5,300 x g for 10 min at 4°C to remove cellular debris. The supernatant was then centrifuged at 21,000 x g for 1 hr at 4°C. Viral RNA was extracted from the viral pellet as previously described [120] and resuspended in 40 μl of RNase-free 5 mM Tris-HCl, pH 8.0. Five μl each of 10 mM dNTPs and 10 μM oligo dT were added prior to denaturing the RNA at 65°C for 10 min. cDNA was synthesized as previously described [121] except at 50°C for 1 hr followed by 55°C for another hour. Two μl RNase H (2U/reaction) were added and the RNA was digested at 37°C for 20 min.

### Single-genome sequencing

To obtain integrase PCR products from single cDNA molecules, cDNA was serially diluted to an endpoint (when approximately 30% of reactions yield PCR product). PCR master mix was prepared as previously described [121, 122] except primers targeting HIV integrase were used: Poli5 (OF) and Poli8 (OR) followed by Poli7 (IF) and Poli6 (IR) [123] in a nested PCR. PCR cycling conditions were modified by decreasing annealing temperature to 50°C and increasing elongation time to 1 min and 30 s. To obtain *env* PCR products from single cDNA molecules, the same diluted cDNA was used but with the following primers, modified from [124]: HIVC.short.VIF1.F1 (5’-GTTTATTACAGGGACAGCAGA-3’) and HIV.short OFM.R1 (5’-CAAGGCAAGCTTTATTGAGGCTTA-3’). Alternatively, E0 forward primer [125] was used. PCR cycling was performed as follows: 95°C for 2 min and 44 cycles of 95°C for 30 s, 57°C for 30 s and 68°C for 3 min – 3 min 30 s, followed by final elongation of 68°C for 5 min. Nested PCR was performed with the following primers, modified from [126]: HIVC.short.ENVA.F2 (5’-GCATCTCCTATGGCAGGAAG-3’) and HIV.short.ENVN.R2 (5’-CAATCAGGGAAGTAGCCTTG-3’). Alternatively, E00 forward primer [125] was used. PCR cycling was performed as follows: 95°C for 2 min, 10 cycles 95°C for 15 s, 57°C for 30 s and 68°C for 3 min, followed by 30 cycles of 95°C for 15 s, 57°C for 30 s and 68°C for 2 min + 5 s per cycle, and a final elongation of 68°C for 7 min. Positive wells were identified by detection using 1% agarose E-Gel^TM^ 96-well gels (ThermoFisher Scientific) and positive PCR reactions were sequenced by Sanger sequencing. For sequencing integrase: Poli7, Poli9D, and O-Poli2C forward and Poli6, Poli10B and Poli3 reverse primers were used [123]. For sequencing *env*: E00, For14, For15, For17, For18 and For19 forward and HIVCshort.ENVN.R2, Rev14, Rev16, Rev17, and Rev18 reverse primers were used [124, 125, 127]. Additional primers used for *env* sequencing are as follows:

For13: 5’-GAGAAAGAGCAGAAGACAGTGG-3’
For16: 5’-TTTAATTGTGGAGGAGAATTTTTCTA-3’
Rev19: 5’-ACTTTTTGACCACTTGCCACCCAT-3’
HIVC.REV20.SEQ: 5’-CACATGGAATTAAGCCAG-3’

### Sequence Analysis

Sanger sequencing data were analyzed using MEGA [128] to align and translate sequences. Sequences were analyzed for known INSTI resistance mutations using HIV Stanford HIV Drug Resistance database: https://hivdb.stanford.edu/

### Statistical analyses

Two-tailed unpaired *t*-tests and Fischer’s exact tests were performed using GraphPad Prism 8 (Graphpad). \**p* < 0.05, ** *p* < 0.01, *** *p* < 0.001 were considered statistically significant. GraphPad Prism was also used to assess statistical significance by one-way ANOVA and Tukey’s multiple comparison test.

## Acknowledgments

We thank members of the Freed laboratory for helpful discussion and critical review of the manuscript. We thank James Thomas (Leidos) for assistance with flow-cytometry analysis. We also thank Michael Bale for assistance with sequence analysis and bioinformatics support. We thank James Hoxie for providing the SupT1 huCCR5 cell line and Alan Engelman for providing BI-224436. The following reagents were obtained through the NIH AIDS Reagent Program, Division of AIDS, NIAID, NIH. Efavirenz, Emtricitabine, Nelfinavir, Maraviroc, AMD3100, T-20, Raltegravir from Merck & Company, Inc, SIM.4 from James Hildreth, sCD4 from Progenics, PGT145, PGT121 and PG16 from IAVI, 2G12 from Polymun Scientific, VRC01 from John Mascola, 447-52D from Susan Zolla-Pazner, 17b from James E. Robinson, 10E8 from Mark Connors, F105 from Marshall Posner and Lisa Cavacini, 35O22 from Jinghe Huang and Mark Connors. 16H3 from Barton F. Haynes and Hua-Xin Liao, HIV-Ig from NABI and NHLBI, pBR-NL43-IRES-eGFP-nef^+^ from Frank Kirchhoff, pK3016 from Christina Ochsenbauer and John Kappes.

## Author Contributions

Conceived and designed the experiments: YH EOF. Performed the experiments: YH RVD PP JLG AW. Analyzed the data: YH RVD JLG MFK EOF. Contributed reagents/materials/analysis tools: YH RVD JLG JWM. Wrote the paper: YH JLG MFK EOF.

## Supporting information

**S1 Fig.**
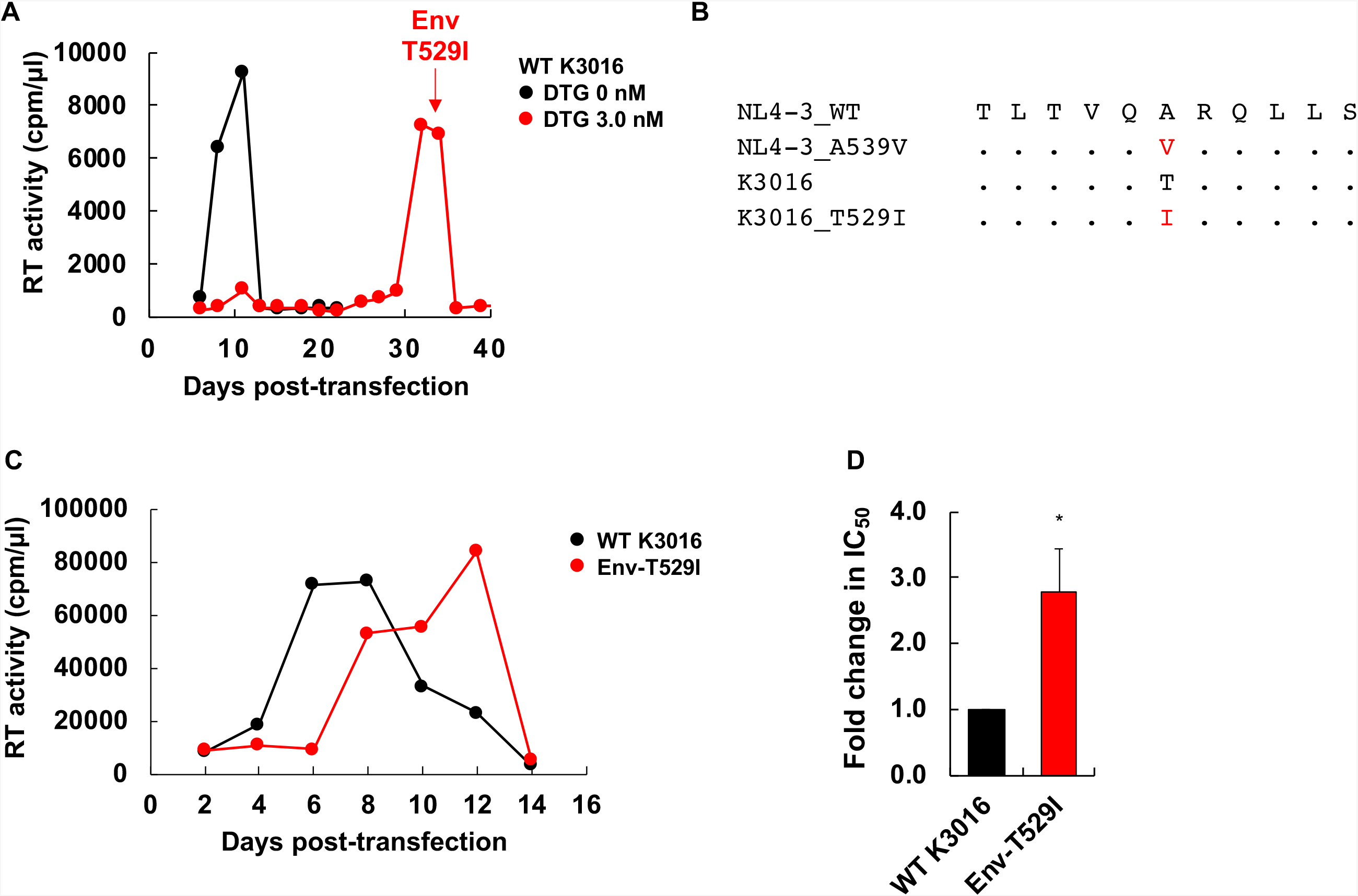
Selection for DTG resistance with the subtype C, CCR5-tropic strain K3016. (A) The SupT1huR5 T-cell line was transfected with pNL(AD8) in the absence or presence of 3.0 nM DTG. At the time point indicated by the arrow, DNA was extracted from the DTG-treated culture and the IN- and Env-coding regions were sequenced, leading to the identification of the Env-T529I mutation. (B) gp41 HR1 sequences around the K3016 Env-T529I mutation are shown aligned with the NL4-3 sequence. (C) The SupT1huR5 T-cell line was transfected with WT K3016 or the Env-T529I derivative in the absence of DTG. The supernatants collected at the indicated time points were assayed for replication kinetics by measuring RT activity. Data are representative of at three independent experiments. (D) The SupT1huR5 T-cell line was transfected with WT K3016 or the Env-T529I derivative in the absence or presence of a serial dilution of DTG (0.03 – 1,000 nM). IC_50_s were calculated based on RT values at the peak of virus replication. Fold changes in IC_50_ relative to WT were calculated. Data from at least three independent experiments are shown as means ± SE. *p*-values < 0.001 (***), < 0.01 (**), < 0.05 (*) by unpaired *t*-test.

**S2 Fig.**
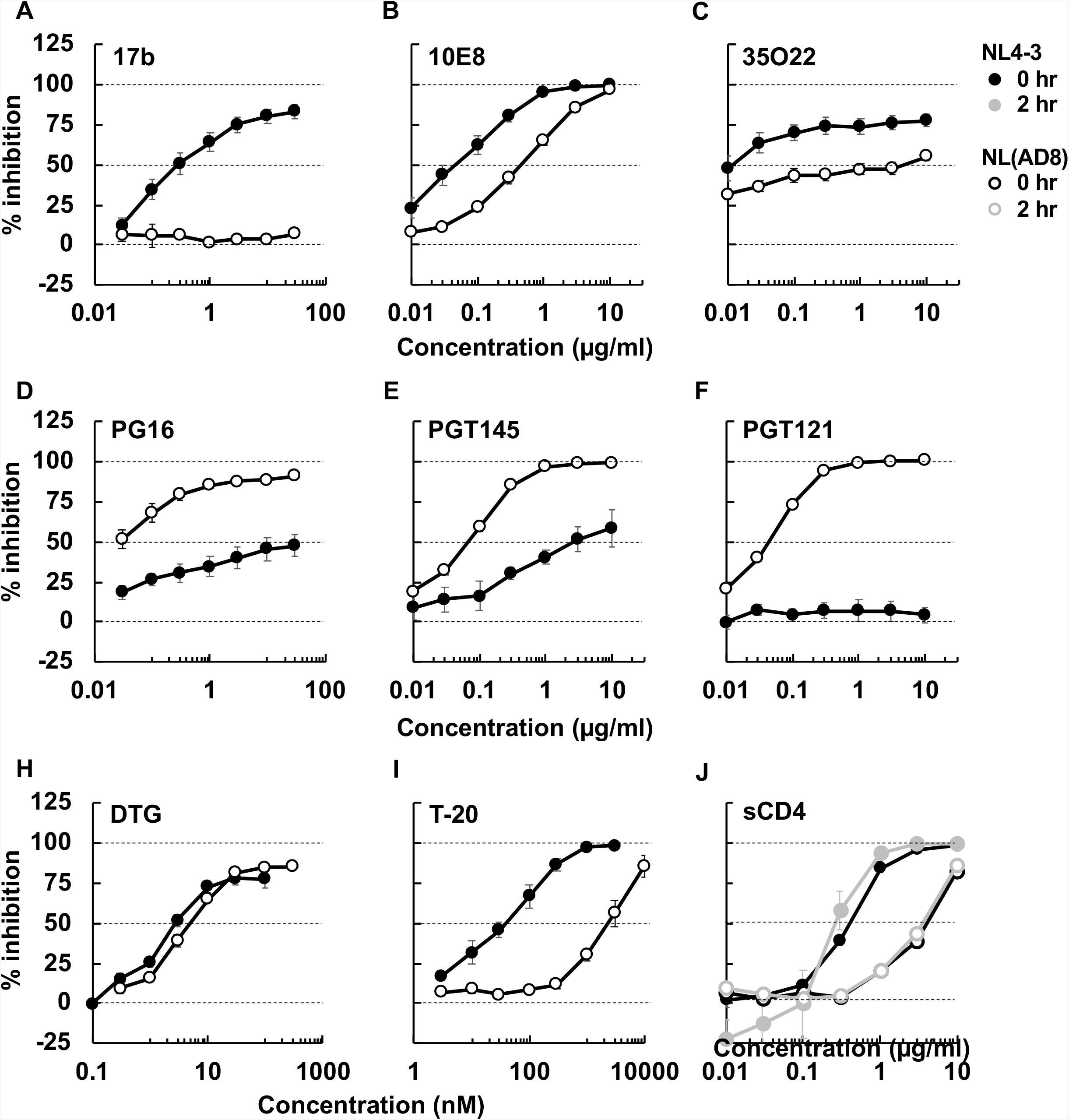
Comparison of the sensitivity of NL4-3 and NL(AD8) to NAbs, DTG, T-20, and sCD4 in cell-free infection. TZM-bl cells were infected with 100 TCID_50_ of the indicated viruses in the presence of varying concentrations of NAbs (A – F), ARVs (H and I), and sCD4 (J). sCD4 incubations were performed for 0 or 2 hrs. The data are related to Fig. 3 and 8. Data from at least three independent experiments are shown as means ± SE.

**S3 Fig.**
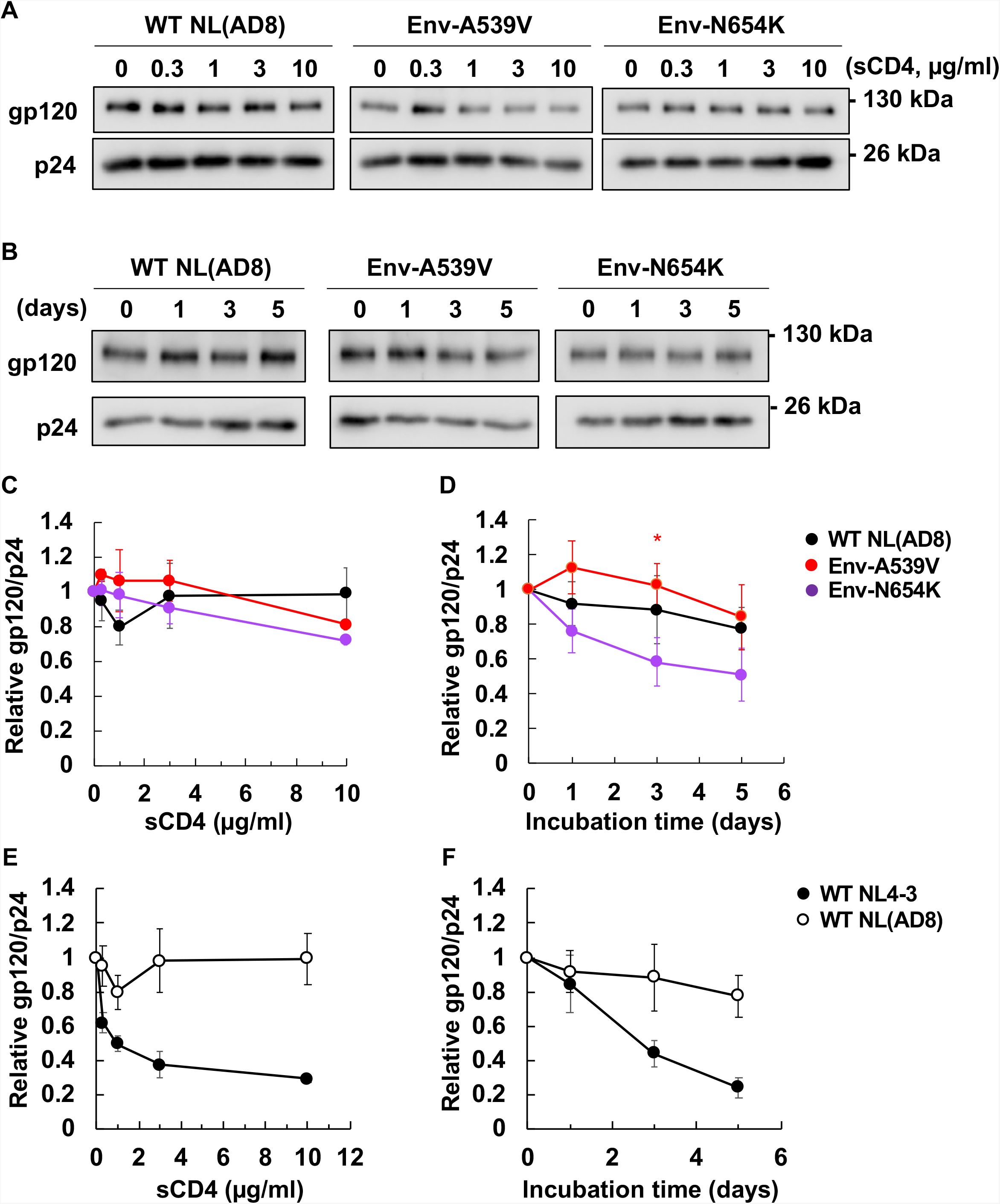
Analysis of sCD4-induced and time-dependent gp120 shedding of NL(AD8) Env mutants and comparison of NL4-3 vs. NL(AD8) gp120 shedding. (A) Concentrated viruses were incubated with the indicated concentrations of sCD4 at 37°C for 2 hrs. Incubated viruses were subsequently purified through 20% sucrose cushions, and viral proteins were detected by western blotting. A representative gel is shown. (B) Viruses were incubated at 37°C for the indicated times. Incubated viruses were subsequently purified through 20% sucrose cushions, and viral proteins were detected by western blotting. A representative gel is shown. Quantification of sCD4-induced (C) and time-dependent (D) gp120 shedding from three independent experiments, calculated as the ratio of virion-associated gp120/p24 and shown as mean ± SE. Comparison of NL4-3 and NL(AD8) sCD4-induced (E) and time-dependent (F) gp120 shedding. The data are related to Fig. 5 and S3. Data from at least three independent experiments are shown as mean ± SE. *p*-values < 0.001 (***), < 0.01 (**), < 0.05 (*), by unpaired *t*-test.

**S4 Fig.**
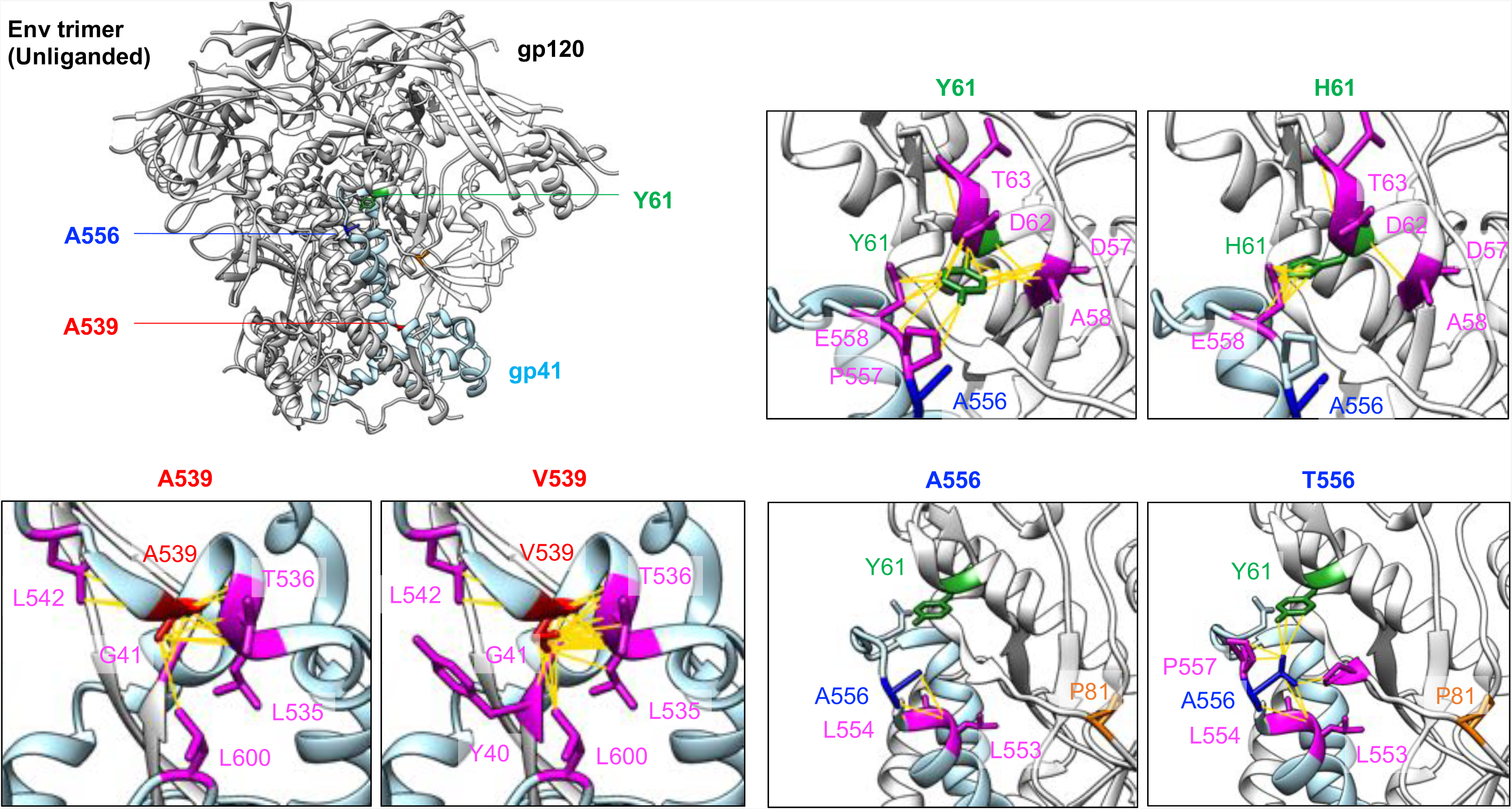
Structural models of NL4-3 Env-Y61H, A539V and A556T mutations. Unliganded Env structure (PDB: 5FYK [96]) in shown at the top left. gp120 and gp41 of protomer 1 are indicated in white and light blue, respectively. Other protomers are indicated in gray. Residues Y61, A539 and A556 are highlighted in green, red and blue, respectively. *In silico* mutagenesis and analysis of possible contacts were performed with UCSF Chimera software [129, 130]. Residues that contact the mutated residues are indicated in magenta. Env amino acid positions are indicated using the NL4-3 numbering. Possible contacts are highlighted as yellow lines.

**S5 Fig.**
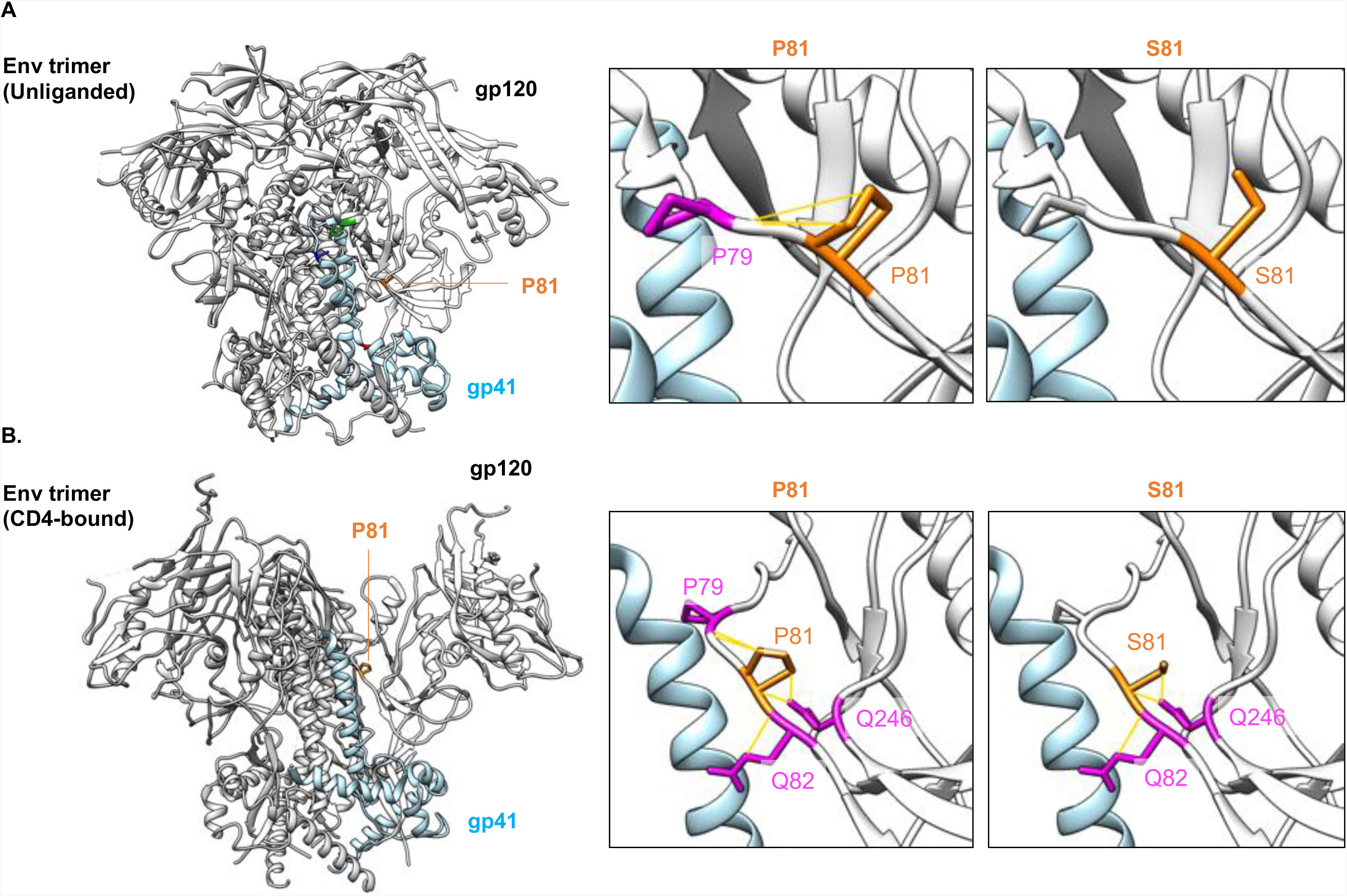
Structural models of the NL4-3 Env-P81S mutant. (A) Unliganded Env structure (PDB: 5FYK [96]) (B) CD4-bound Env structure (PDB: 5VN3 [95]). gp120 and gp41 of protomer 1 are indicated in white and light blue, respectively. Other protomers are indicated in gray. Residue P81 is highlighted in orange. *In silico* mutagenesis and analysis of possible contacts were performed by UCSF Chimera software [129, 130]. Residues that contact the mutated residues are indicated in magenta. Env amino acid positions are indicated using the NL4-3 numbering. Possible contacts are highlighted as yellow lines.

